# Influence of duty-cycle recording on measuring bat activity in passive acoustic monitoring

**DOI:** 10.1101/2025.04.15.649046

**Authors:** Aditya Krishna, Wu-Jung Lee

**Affiliations:** Applied Physics Laboratory, University of Washington, Seattle, Washington, USA; Department of Electrical and Computer Engineering, University of Washington, Seattle, Washington, USA

## Abstract

Echolocating bats provide vital ecosystem services and can be monitored effectively using passive acoustic monitoring (PAM) techniques. Duty-cycle subsampling is widely used to collect PAM data at regular ON/OFF cycles to circumvent battery and storage capacity constraints for long-term monitoring. However, the impact of duty-cycle subsampling and potential detector errors on estimating bat activity has not been systematically investigated for bats. Here, we simulate the influence of duty-cycle subsampling in measuring bat activity via three metrics–Call Rate (CR), Activity Index (AI), and Bout-Time Percentage (BTP)–using three months of continuous recordings spanning summer to fall in a temperate urban natural area. Our simulations show that subsampled bat activity estimates more accurately track true values when the listening ratio is high and the cycle length is low, when the true call activity is high, or when recorded calls have lower frequency content. Generally, among the three metrics, AI provides the best subsampling estimates and is robust against false negatives but sensitive to false positives, whereas BTP provides better temporal resolution compared to AI and is robust against both false positives and false negatives. Our results offer important insights into selecting sampling parameters and measurement metrics for long-term bat PAM.

## I. INTRODUCTION

Passive acoustic monitoring (PAM) is a powerful technique for monitoring soniferous animals in both terrestrial and marine habitats, and has in recent years been widely adopted thanks to broader availability and accessibility of low-cost recording devices (Gibb et al., 2019; Hill et al., 2018; Lahoz-Monfort and Magrath, 2021; Atkin, 2024). Bats are a good target animal group for PAM because of their nearly ubiquitous use of ultrasonic echolocation signals for foraging and navigation (Moss et al., 2023). As bats’ echolocation signals vary depending on the species and behavioral context (Ratcliffe et al., 2013; Russo et al., 2018), such as prey searching or capture, PAM data are useful in identifying both the spatiotemporal distribution of different species and potential changes in habitat use in response to environmental variabilities—such as broader climate pattern changes—and to the emergence of infectious diseases, such as white-nose syndrome (WNS) (Hicks et al., 2020; Ross et al., 2023).

Duty-cycle subsampling is a sampling method widely used to mitigate the logistical and financial constraints associated with collecting and storing long-term PAM data (Francomano et al., 2021; Melo et al., 2021; Pieretti et al., 2015). Under duty-cycle subsampling, data is collected with regular ON/OFF cycles, trading sampling continuity with increased total deployment duration (Thomisch et al., 2015). Depending on the configuration, subsampling can improve the efficiency of detecting specific species and differentiating the relative abundance of different species (Cook and Hartley, 2018) while retaining soundscape information contained in continuous recordings, such as temporal autocorrelation ranges and variability of acoustic indices (Francomano et al., 2021; Pieretti et al., 2015). For bat monitoring, multiple studies have empirically derived minimal PAM sampling efforts needed to fully describe the species assemblage and its spatial and temporal changes (Baumgardt et al., 2022; Hauer et al., 2023; López-Baucells et al., 2021). However, there remains a need for systematic investigations into the impacts of duty-cycle subsampling on measuring bat activity levels, as such estimates can vary greatly depending on the call characteristics and behavioral context of the target species (Rand et al., 2022; Thomisch et al., 2015).

In addition, the interactions among temporal subsampling configurations, different metrics of bat activity, and detector error characteristics remain unclear. Indeed, the difficulty of mapping the number of detected calls to bat abundance has led to development of many metrics or proxies, including using contextual cues (Hoggatt et al., 2024; Milchram et al., 2020), metrics based on call intensity (Kloepper et al., 2016), and temporally aggregated metrics that count the number of predetermined time blocks with bat call occurrence (activity index) or use bat passes (Beason et al., 2020; Miller, 2001; Tuneu-Corral et al., 2020). As automatic call detection algorithms become essential for managing the ever-growing volumes of PAM data, understanding the potential impacts of detector biases, including false detections (false positives) and misses (false negatives) (Gibb et al., 2019), is especially crucial for accurate data interpretation.

To address these gaps, in this paper we investigate the influence of duty-cycle subsampling in measuring bat activity, focused on two core questions: 1) How does the configuration of duty-cycle subsampling affect bat activity estimates compared to true values from continuous data under different bat activity levels and call characteristics? and 2) How do different activity metrics perform in response to duty-cycle subsampling and detector error? We used a PAM dataset collected continuously (24/7) from a temperate urban natural area that experienced diverse environmental fluctuations in temperature and precipitation. Using this dataset, we simulated a wide range of subsampling configurations and evaluated the performance of three activity metrics against true values derived from the continuous data. Our goal is to enhance the accuracy and efficiency of PAM in bat monitoring by providing guidelines for selecting effective duty-cycle subsampling schemes and activity metrics depending on call activity and call characteristics.

## II. METHODS

### A. Data collection

Passive acoustic recordings were collected from six sites in the Union Bay Natural Area, an urban natural area adjacent to the University of Washington’s Seattle campus (Fig. 1). The sites range from open ponds to areas under dense canopy, attracting different dominant bat species, such as big brown bats (*Eptesicus fuscus*) and little brown bats (*Myotis lucifugus*). This provided us with a broad range of bat activity levels and call characteristics to systematically investigate the influence of duty-cycle subsampling configuration under a variety of monitoring conditions.

**FIG. 1.**
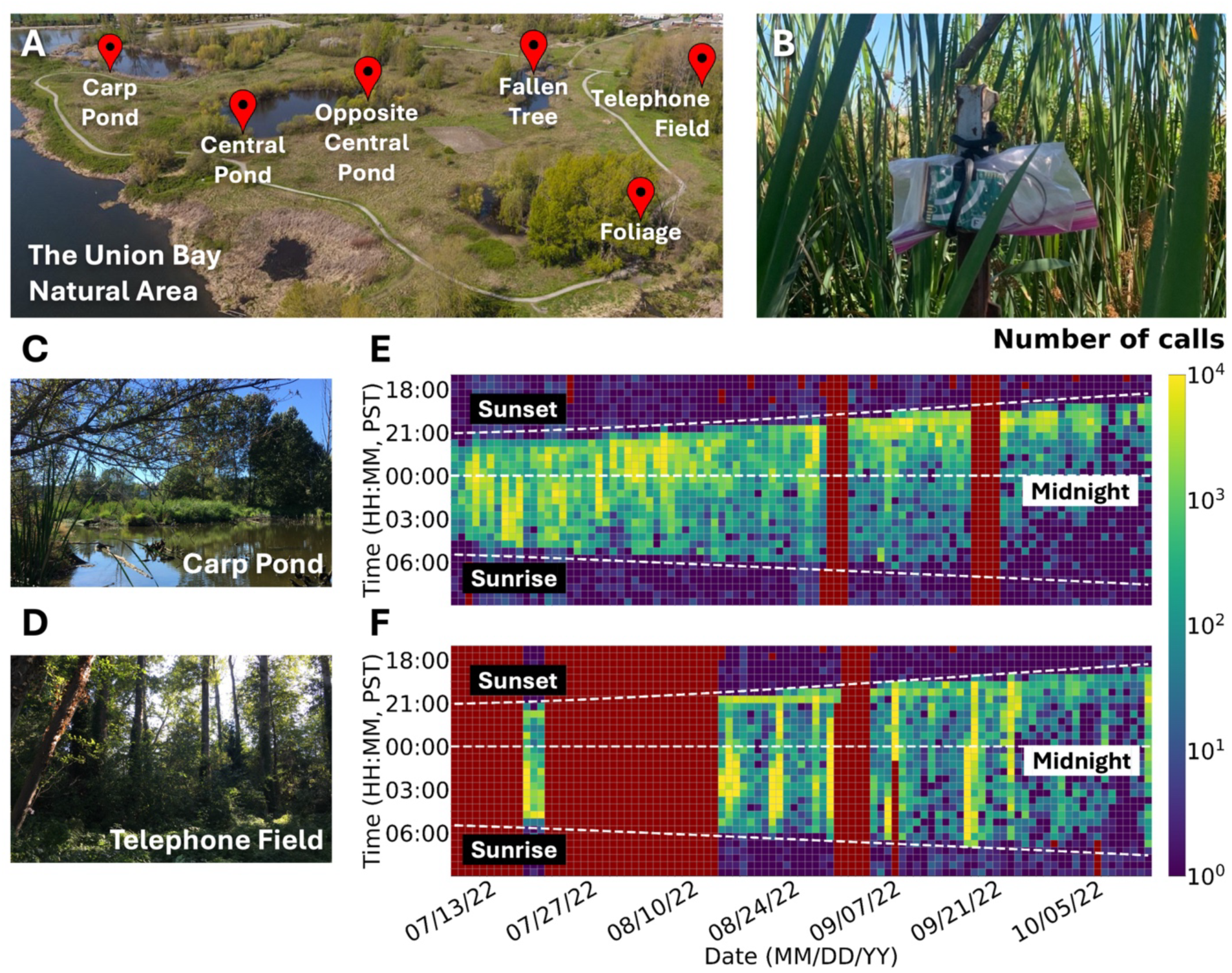
(A) Recording sites in the Union Bay Natural Area. (B) An AudioMoth used for passive acoustic recording. (C) A site next to an open pond (“Carp Pond”). (D) A site under a dense canopy (“Telephone Field”). (E-F) Example data overview showing the number of bat echolocation calls detected in 30-min windows at Carp Pond (E) and Telephone Field (F) during the data time period used in this study. Dark red indicates periods in which no data was recorded.

The recordings were collected from July to October 2022 using the AudioMoth v1.2.0 recorders (Hill et al., 2018) placed inside plastic Ziploc bags and attached to a tree branch or a metal stake (Fig. 1B) approximately 1.5-2 m above the ground facing an open space. Each recorder was equipped with either three or six 2450mAh NiMH AA batteries and 128GB microSD cards. The recorders were configured to continuously record data at a gain setting of medium and a sampling rate of 250 kHz until late July. The sampling rate was switched to 192 kHz once we confirmed that no recorded bat calls had significant frequency content between 96 and 125 kHz.

### B. Call detection and classification

We used two automated detectors, the open-source `batdetect2` (Aodha et al., 2022) and the proprietary Kaleidoscope Pro 5.6.8 (Wildlife Acoustics, Maynard, Massachusetts), to extract bat search-phase calls. Raw `batdetect2` detections were filtered using thresholds based on the detection confidence score and signal-to-noise ratio (SNR) of the calls. These thresholds were tuned by evaluating detector performance against a set of human-annotated calls (see Supplemental Materials Section S1). We performed all analyses using both `batdetect2` and Kaleidoscope Pro, but present only results based on the former in the main text due to its better performance (Table S1). We performed analyses using calls detected from July to October, 2022, to focus sampling during the pre-hibernal period (Schowalter, 1980; Speakman and Rowland, 1999).

In this study, we classified all detected bat calls into two “phonic groups”: low-frequency (LF) and high-frequency (HF), to analyze based on calls with similar frequency content (Fraser et al., 2020; Ober and Hayes, 2008), instead of species. This is because: 1) in the context of PAM, call frequency content plays an important role in determining call detectability, as bat ultrasonic echolocation calls are subject to strong atmospheric absorption that intensifies with increasing frequency (Stilz and Schnitzler, 2012); and 2) the uncertainty of identifying species using echolocation calls alone, especially for species in the genus *Myotis* whose call spectra and frequency contour overlap significantly (Russo et al., 2018). The LF group (25-40 kHz) likely contained *Eptesicus fuscus*, *Lasionycteris noctivagans*, and *Lasiurus cinereus*, and the HF group (35-80 kHz) likely contained *Myotis lucifugus, Myotis yumanensis, and Myotis californicus* based on Kaleidoscope Pro’s “Bats of North America 5.4.0” classifier filtered for the Washington region, under the neutral setting. Calls were classified into the LF and HF phonic groups (Fig. 2) using a *k*-means classifier trained on a representative set of high-quality call samples selected from all recording files (see next paragraph for details). We determined the number of clusters (*k*=2) based on a preliminary visual inspection of the detected call spectra, along with hierarchical agglomerative clustering results (using average linkage and Euclidean distance), which confirmed the existence of two main phonic groups. Both the agglomerative clustering and *k*-means clustering were based on the Welch power spectral densities (PSD) of the calls, with each call normalized by its spectral maximum and interpolated to a standard frequency vector spanning 0-96 kHz. Misclassified bat calls from the *k*-means classifier were identified based on the call peak frequency and removed for each 30-min recording segment (see Supplemental Materials Section S2). Out of all 4,801,832 detected calls used in the analysis, 68,670 were misclassified by the *k*-means classifier.

**FIG. 2.**
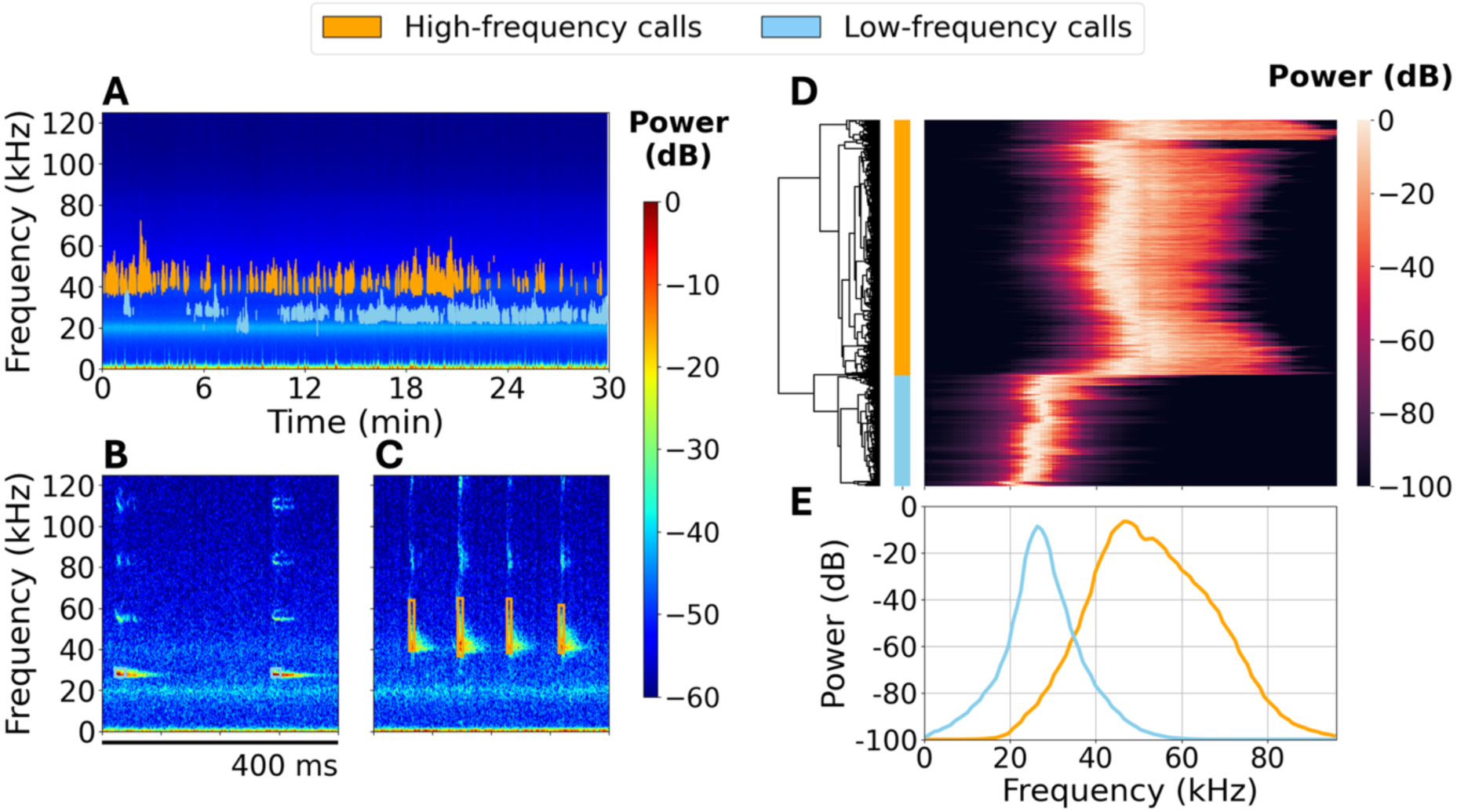
Classification of phonic groups. (A) Spectrogram of a 30-min recording with detected bat calls classified into either HF or LF phonic groups. (B-C) Example calls from the HF and LF phonic groups. Call bounding boxes plotted are time and frequency extents given by `batdetect2`. (D) Dendrogram of normalized power spectral density (PSD) for all calls in the representative set (see Sec. II. B for detail). (E) Centroid PSD derived from *k*-means clustering of representative call samples. Calls in panels B-D are color-coded according to their *k*-means cluster membership in panel E.

The representative set of high-quality bat call samples was derived from sites: Carp Pond, Central Pond, Foliage, and Telephone Field. Call samples were selected using the following procedure. First, calls detected by `batdetect2` with a confidence score ≥0.5 were assigned into three crude frequency bands (13-50 kHz, 30-78 kHz, or 41-96 kHz) based on the high- and low-frequency extents given by `batdetect2`. These crude bands were selected based on visual inspection. Within each band, calls were grouped into “bouts,” or sequences of calls with small spacings (see Sec. II. C for detail). ‘Clipped’ echolocation calls (calls that saturated the recorder electronics and led to distorted waveforms) were present in our data. However, because our classification procedure was based on the broad spectral content of the calls rather than fine spectral features, the influence of clipped calls was likely minimal. The representative set was then assembled by selecting calls with the top 1% SNR within each bout. Since different bouts varied significantly in their overall SNR, this procedure ensured that low-SNR calls were also included in the representative set. Here, SNR was defined as the ratio between the total power of a bat call and the total power of the “noise” segment of equal duration immediately preceding the call. Both call and noise segments were bandpass filtered using a 4th-order Butterworth filter with cutoff frequencies set at 2 kHz below the low-frequency extent and 2 kHz above the high-frequency extent given by `batdetect2`. Note that the initial crude frequency bands mentioned here do not play a role in the downstream clustering analyses.

### C. Metrics of activity

Acoustics-based metrics, such as call rate (CR) and activity index (AI), have been widely used to characterize temporal fluctuations of bat activity (Brinkløv et al., 2023; Skalak et al., 2012). In this study, we used CR, AI, and bout-time percentage (BTP), a metric we developed using the concept of “bouts”, to characterize bat activity. These metrics are defined below.

#### Call rate (CR, Fig. 3A)

CR is the number of calls within a specific time duration. It is easy to compute and is an absolute and continuous measure. In this study, we compute the call rate in terms of the number of calls per minute.

#### Activity index (AI, Fig. 3B)

AI is computed by dividing a given time period into smaller time blocks and counting the number or percentage of blocks containing at least 1 call within a given time period. This metric was proposed by (Miller, 2001) and often used for cross-species activity comparisons, as it minimizes the influence of inherent differences in call rates (Froidevaux et al., 2014; Skalak et al., 2012). In this study, we use a time block size of 5 seconds to detect differences in activity with fine temporal resolution (Miller, 2001).

#### Bout-time percentage (BTP, Fig. 3C)

BTP is computed by first grouping calls into “bouts” and then calculating the percentage of time occupied by all bouts within a given time period. In animal behavior literature, bouts are defined as repeating events of interest that are grouped together when the interval between events is less than the bout criterion interval (BCI) (Fagen and Young., 1978; Sibly et al., 1990; Slater and Lester, 1982). BCI models the between-event intervals as generated from a Poisson process characterized by either the between-bout interval (BBI) or the within-bout interval (WBI) based on a log-survivorship curve. In this study, we define each call as an event and use the inter-pulse interval (IPI) as the between-event interval. Among the three methods proposed to find the BCI (Fagen and Young., 1978; Sibly et al., 1990; Slater and Lester, 1982), we chose to follow the Slater-Lester method for its simplicity and reliable BCI estimates (see Supplemental Materials Section S3). Unlike AI that requires a fixed time block size, BTP is a continuous measure which measures the percentage of time bat calls were detected as bouts within a given time period.

### D. Simulation and evaluation of duty-cycle subsampling

To investigate the influence of duty-cycle subsampling on measuring bat activity, we simulated subsampling by systematically varying the cycle length (the sum of recorder ON and OFF time per cycle) across 6, 10, 30, 60 minutes and the listening ratio (the proportion of the cycle when the recorder is ON) across ⅙, ⅓, ½, ⅔. This resulted in a total of 16 subsampling schemes, defined as combinations of cycle length and listening ratio. Cycle length and listening ratio are two independent parameters that can be used to define duty-cycle subsampling schemes (Fig. 3D). We simulated subsampled data by dividing the continuous data into consecutive, non-overlapping time periods based on the cycle length and removing the calls falling within the OFF period. We then treat each time period as a sample that can be used to evaluate the quality of activity metric derived from subsampled data by comparing the metric value to its continuous counterpart. To quantitatively compare the effect of subsampling for each activity metric, we calculated the percentage of duty-cycle samples for which the estimated activity metric values fall within an error bound around the true activity metric values (Fig. 4).

**FIG. 3.**
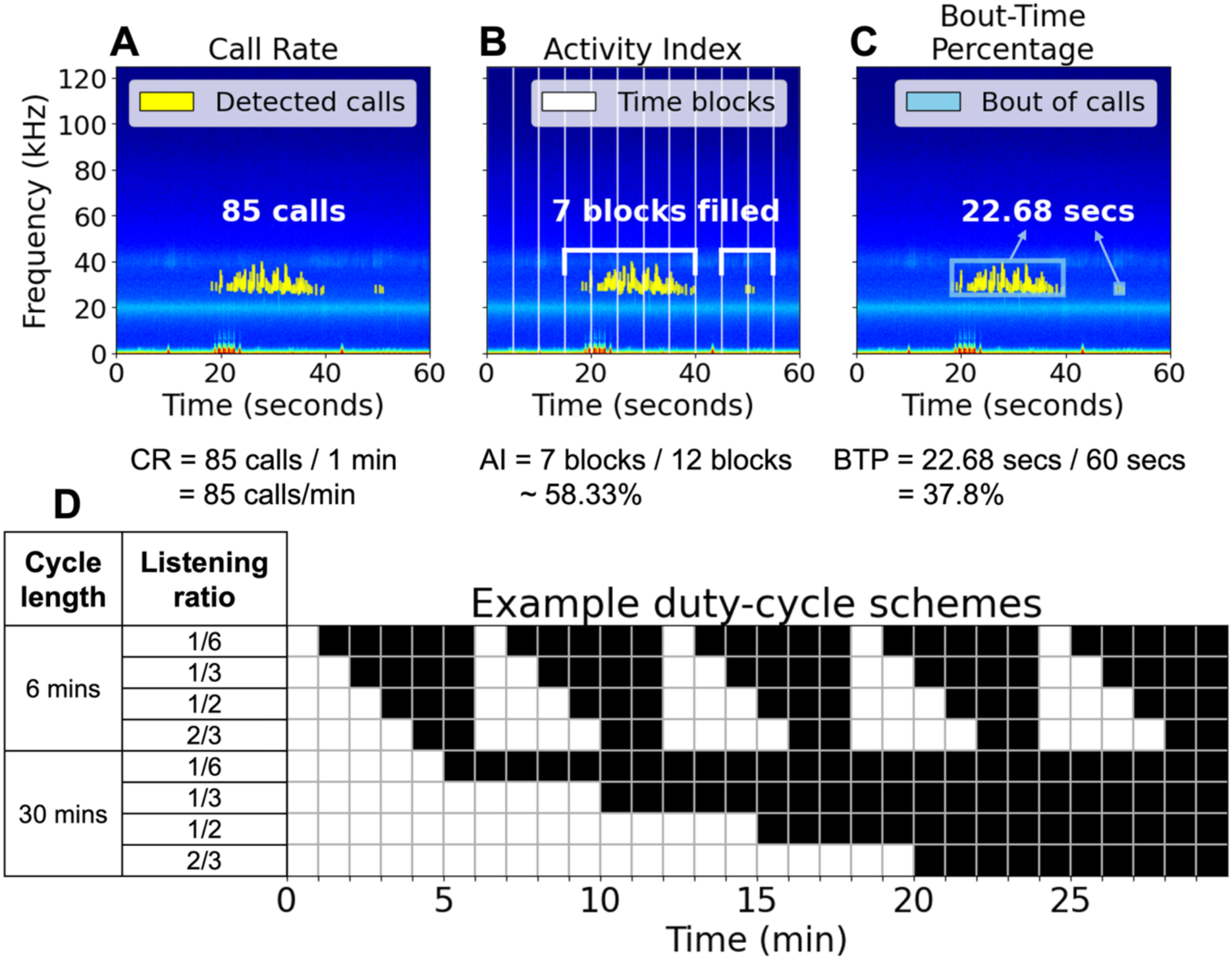
An example 1-minute recording segment showing the calculation of the three activity metrics and a diagram showing example duty-cycle schemes. (A) Call rate (CR) is the average number of calls per minute in a given recording segment. (B) Activity index (AI) is the percentage of 5-sec blocks occupied with calls within a given recording segment. (C) Bout-time percentage (BTP) is the percentage of time occupied by bouts within a given recording segment. (D) Example duty-cycle schemes for a 30-min recording period with cycle lengths of 6 and 30 mins and varying listening ratios (see Sec. II. D for definition). White blocks indicate recorder ON times and black blocks indicate recorder OFF times.

**FIG. 4.**
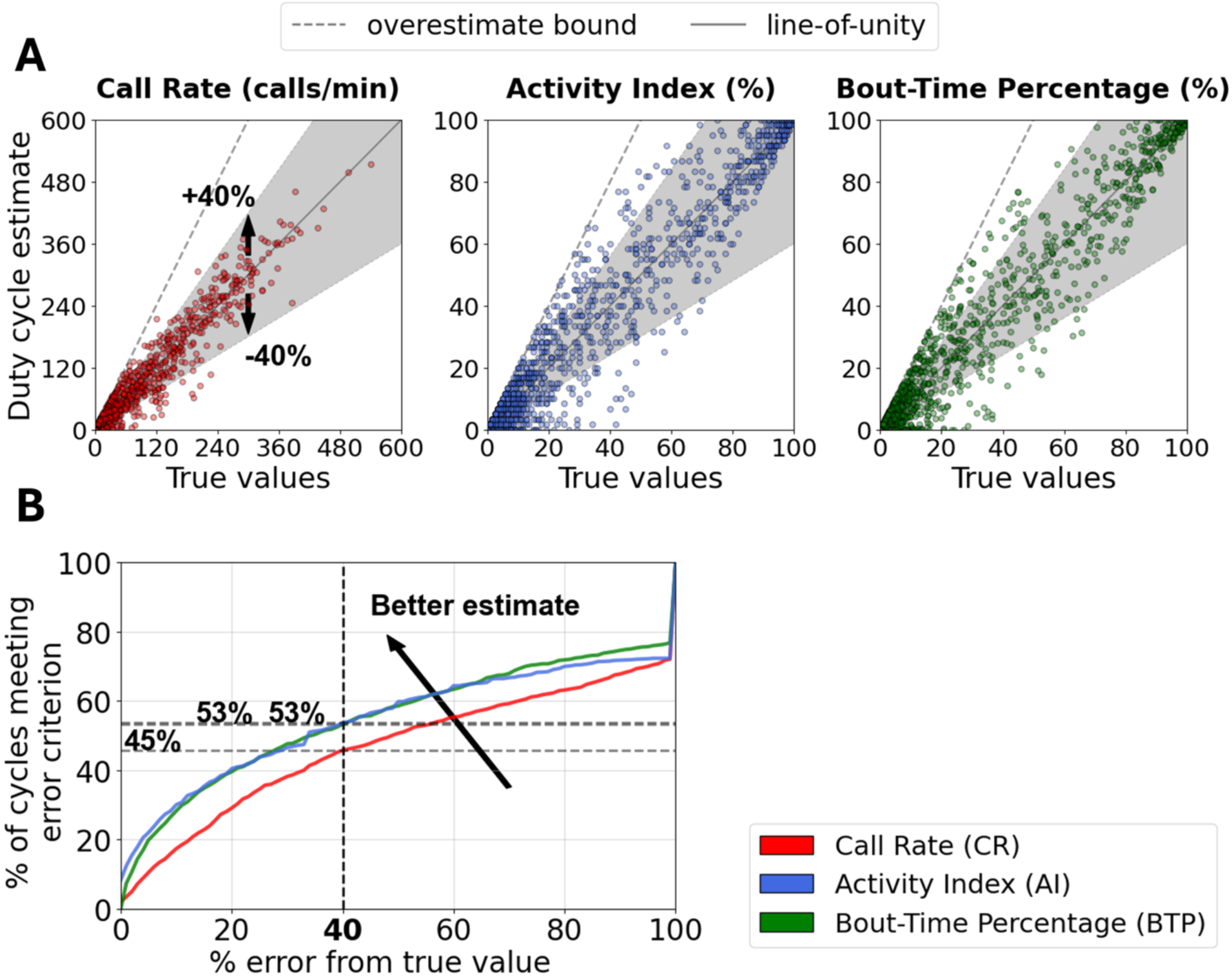
The procedure to generate an evaluation curve showing the quality of activity metric estimates. (A) Comparison of subsampling estimates and true values. Each point represents a duty-cycle sample. The slanted grey dashed line is the overestimate bound (see Supplemental Materials Section S4). The grey region indicates the area in which the duty-cycle samples fall within an error bound (set at 40% in this example). (B) Evaluation curves showing the percentage of duty-cycle samples meeting the error criterion across increasing error bounds for each activity metric. Samples meeting the error criterion are located within the specified error bounds (i.e., within the grey region) in panel A. A faster rise in the evaluation curve indicates a higher percentage of duty-cycle samples appearing near the line-of-unity and therefore better quality estimates. All panels were generated using duty-cycle samples of Carp Pond LF calls during July-August, 2022, with a cycle length of 10 mins and a listening ratio of 1/2.

We analyzed the influence of subsampling on all three activity metrics given different overall bat activity levels, phonic groups, and detector error characteristics in terms of false positives and false negatives (misses). For overall bat activity, we categorized duty-cycle samples into high activity and low activity conditions using thresholds of CR≥50 calls/min, AI≥50%, and BTP≥50% measured in the continuous dataset. For phonic groups, we compared the subsampling effects of the LF phonic group recorded at Carp Pond with the HF phonic group recorded at Telephone Field, because these datasets exhibited similarly high overall activity levels. For detector error characteristics, we treated detections from `batdetect2` as a proxy for “true” values and simulated detectors with a *p*% false positive or false negative error rate for high and low activity conditions separately. These detections are proxies because automated detectors have inherent errors and cannot provide perfect ground truth. False positives were generated by randomly inserting *N*_*false*_ artificial 10-msec detections into the original set of detections for each 30-min recording segment. False negatives were generated by removing *N*_*false*_ detections from the original recording segments. Here, *N*_*false*_ = *floor*(*p*% × *N*_*T*_), where *N*_*T*_ is the total number of originally detected calls in a given recording segment. For 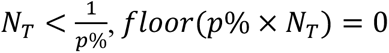. Hence, for these cases it is expected that *N*_*false*_ = 0 and there will be no difference in the simulated and true estimates.

## III. RESULTS

### A. Continuous dataset

During the 2022 recording period, a total of 5453 hours of audio were collected from the four sites considered representative (see Sec. II. D). The number of calls, bout duration, and BBI varied widely across sites. Among the sites, Carp Pond and Telephone Field had the highest number of recorded LF and HF calls, respectively. Carp Pond LF calls and Telephone Field HF calls differed in call peak frequencies, but both groups showed high overall activity and were chosen for subsequent analyses (Fig. 5). Groups such as Carp Pond HF, Telephone Field LF, Foliage LF and HF, and Central Pond LF and HF calls were not used in subsequent analyses because they exhibited lower levels of overall activity, which made it difficult to perform analyses requiring periods of varying activity levels. Only recordings from 03:00 to 13:00 UTC (20:00 to 06:00 PST; local night time) were used.

**FIG. 5.**
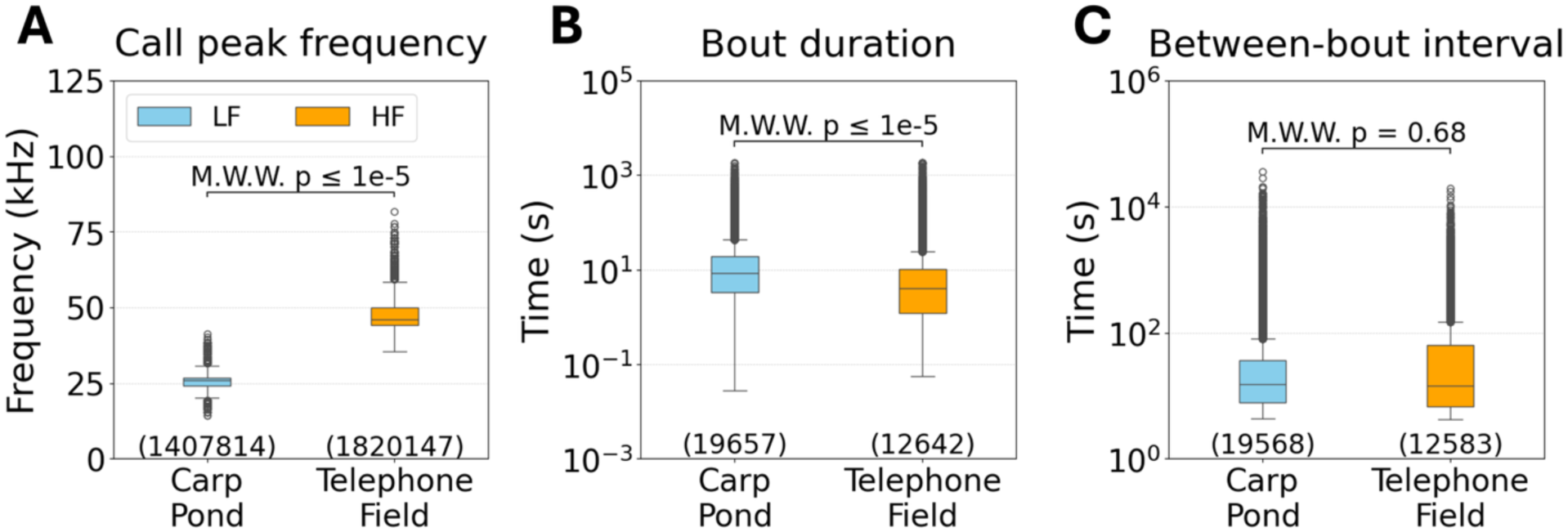
Call and bout parameters from Carp Pond LF calls and Telephone Field HF calls. (A) Call peak frequency. (B) Bout duration. (C) Between-bout interval (BBI). The number of samples included are shown at the bottom of each boxplot. In each boxplot, the central line represents the median, the box spans the interquartile range (IQR, 25th to 75th percentile), and the whiskers extend to the minimum and maximum values within 1.5 times of IQR. Outliers beyond this range are shown as individual points. M.W.W.: Mann-Whitney-Wilcoxon test.

### B. Duty-cycle configuration

Our simulations showed that increasing listening ratios under a fixed cycle length provides better subsampling estimates for all activity metrics (Fig. 6A, Fig. S12, Fig. S13). For example, with a fixed cycle length of 30 mins, increasing the listening ratio from ⅙ to ½ improves the estimates of all three activity metrics (Fig. 6A). Separately, decreasing the cycle length while keeping a fixed listening ratio provides better subsampling estimates for all activity metrics (Fig. 6B, Fig. S12, Fig. S13). For example, with a fixed listening ratio of ½, decreasing the cycle length from 60 mins to 10 mins improves the estimates of all three activity metrics (Fig. 6B).

**FIG. 6.**
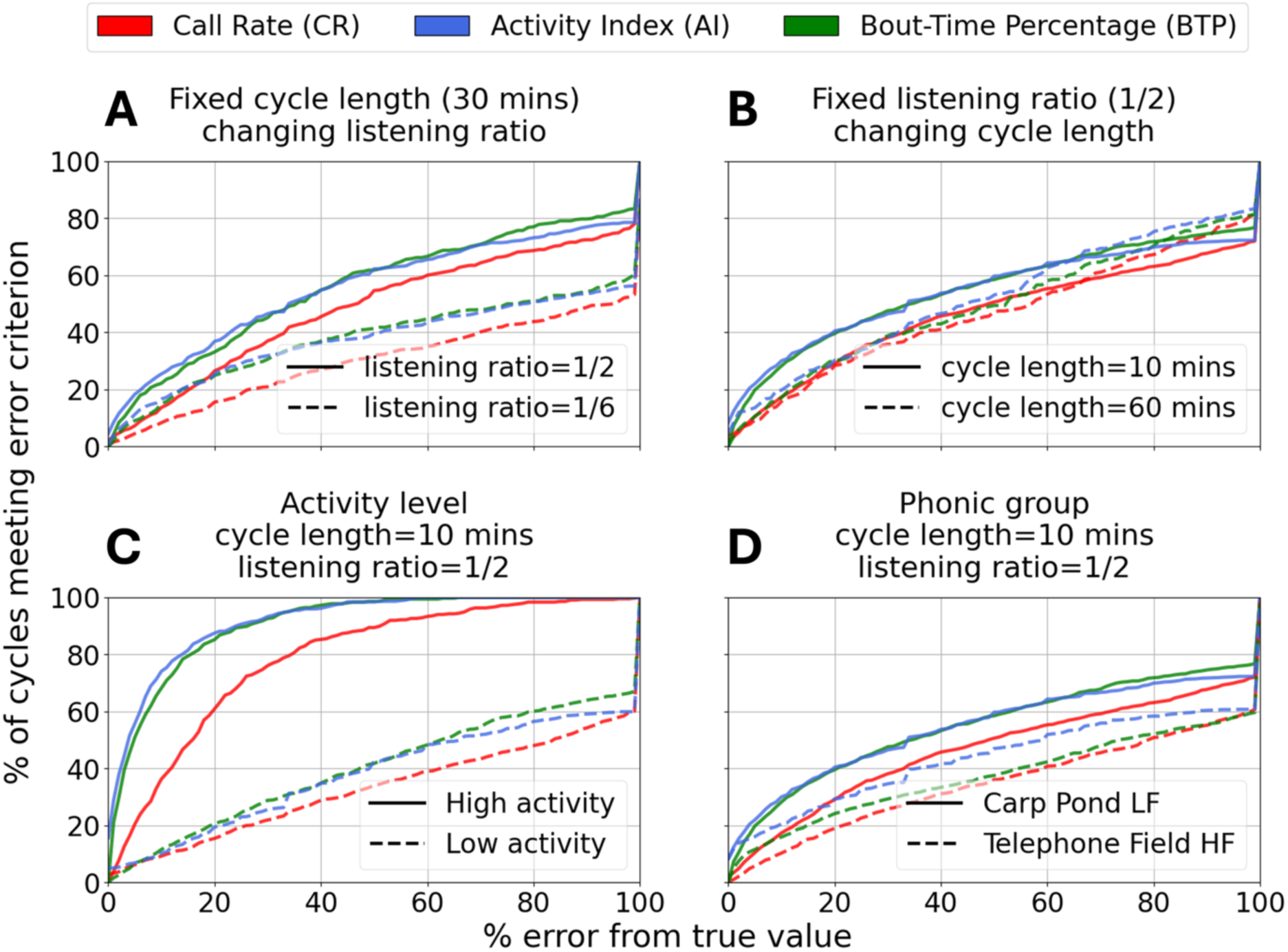
Quality of duty-cycle subsampling estimates. (A) Influence of listening ratio with a fixed cycle length. (B) Influence of different cycle lengths with a fixed listening ratio. (C) Influence of overall activity level on duty-cycle estimates. (D) Influence of call frequency content on duty-cycle estimates. LF (solid) calls from Carp Pond and HF (dashed) calls from Telephone Field were separately evaluated with the same duty-cycle scheme and compared. Panels A-C were generated using LF calls recorded at Carp Pond during July-August, 2022. In panel D, the LF calls used came from the same period as in panel A-C, while the HF calls came from Telephone Field recordings during August-September, 2022.

### C. Activity level and phonic groups

Under different overall activity levels (see Sec. II. D for definitions), we found that duty-cycle subsampling estimates are more accurate for high activity periods than low activity periods (Fig. 6C, Fig. S14, Fig. S15). For example, with a listening ratio of ½ and a cycle length of 10 mins (Fig. 6C), duty-cycle estimates from periods of high activity significantly outperform the estimates from periods of low activity for all three activity metrics.

Across the two phonic groups with substantially different call frequency content, our simulation showed that duty-cycle estimates for LF call activities outperform the estimates for HF call activities for all three activity metrics (Fig. 6D, Fig. S16).

### D. Activity metrics

Across all duty-cycle configurations, activity levels, and phonic groups, subsampling estimates of AI and BTP often track one another closely, with both consistently outperforming estimates of CR (Fig. 6, Fig. S12-16). Below we further evaluate the robustness of these activity metrics against detector errors in terms of false positives (false detections) and false negatives (misses) across high and low activity levels as categorized in Sec. II. D.

For continuous data, adding false positives appeared to impact AI and BTP estimates more than adding false negatives (Fig. 7A and 7D). For low activity periods, all three activity metrics included duty-cycle samples that had 0% error from true value (i.e., non-zero y-axis intercepts) when no simulated false positive or false negative calls were added (*N*_*false*_ = 0, see Sec. II. D). Duty-cycle estimates of BTP are consistently better than those of AI when false positives are present (Fig. 7A), whereas duty-cycle estimates of AI and BTP closely track each other when false negatives are present (Fig. 7D). For high activity periods, duty-cycle estimates of BTP and AI closely track each other when either false positives or negatives are present (Fig. 7A and 7D). Duty-cycle estimates of CR exhibited a sharp rise from 0 to 100% regardless of activity level when the error tolerance crossed the false positive or false negative rate (Fig. 7A and 7D). This sharp rise is intuitive, since with a fixed percentage of calls added or removed in our simulation (20% in the examples shown), no recording periods would meet the error criterion when error tolerance was less than 20%.

**FIG. 7.**
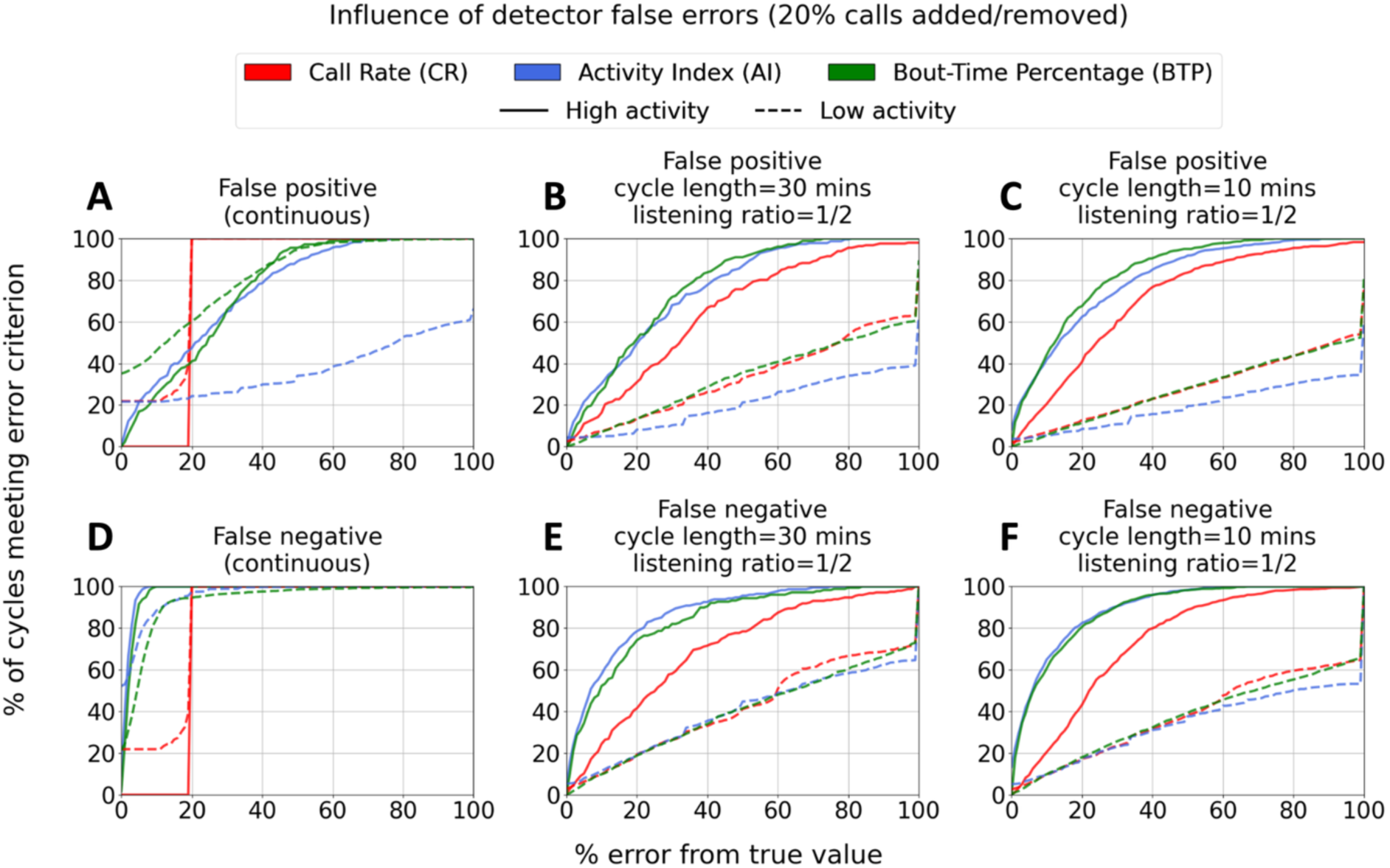
Quality of duty-cycle subsampling estimates during high and low activity periods with simulated false positives and false negatives. (A-C) Influence of false positives. Here, false positives were simulated by randomly adding 20% more 10-msec artificial detections into the data. (D-F) Influence of false negatives. Here, false negatives were simulated by randomly removing 20% of the detected calls from the data. All simulated call addition or removal were performed prior to subsampling and computing activity metrics. See Sec. III. D for details of simulating false positives and false negatives. All panels were generated using LF calls from Carp Pond during July-October, 2022. Continuous: no duty-cycle subsampling.

For subsampled data, the presence of false positives or false negatives impacted all three activity metrics (Fig. 7B-C and 7E-F). For low activity periods, duty-cycle estimates of BTP consistently track those of CR while both consistently outperform estimates of AI when false positives were present (Fig. 7B-C), whereas duty-cycle estimates of all three metrics track one another closely when false negatives were present, especially when the error bounds ≲60% (Fig. 7E-F). For high activity periods, duty-cycle estimates of BTP and AI generally tracked each other and outperformed CR, regardless of whether false positives or negatives were present (Fig. 7B-C and 7E-F).

## IV. DISCUSSION

In this study, we systematically analyzed the influence of duty-cycle subsampling on estimating bat activity level using three metrics—CR, AI, and BTP—that summarize the temporal occurrence of bat calls. We simulated duty cycle recording with varying cycle length and listening ratio to subsample a continuously recorded dataset (the “truth”), and investigate how subsampling impacts estimates of these metrics under different overall call activity and call frequency content, and how different metrics respond to potential detector errors.

Our simulations showed that increasing the listening ratio within a cycle always improved duty-cycle subsampling estimates for all activity metrics, irrespective of the cycle length (Fig. 6A, Fig. S12, Fig. S13). These results are intuitive, as a higher proportion of data is retained with increasing listening ratio. In the extreme case when the listening ratio is 1, all data is retained and no errors would exist between duty-cycle estimates and the true values. Our results are consistent with previous studies that showed increasing listening ratio improved the accuracy of estimating indices such as species richness, behavioral patterns, and soundscape variability for both terrestrial and marine animals (Anunciação et al., 2022; Francomano et al., 2021; Pieretti et al., 2015; Rand et al., 2022; Riera et al., 2013). However, as also discussed previously, a higher listening ratio increases power consumption and storage requirements, making it an important trade-off that depends on available resources and cost.

We also found that decreasing cycle length always improved duty-cycle subsampling estimates for all activity metrics, irrespective of the listening ratio (Fig. 6B, Fig. S12, Fig. S13). This can be intuitively understood by considering the stationarity of call activity. Duty-cycle subsampling assumes that the monitored quantity—be it call activity or diversity—during the recorder ON period accurately represents the same quantity during the recorder OFF period. Longer cycle lengths result in longer recorder OFF periods, making this assumption increasingly tenuous and leading to a higher likelihood of subsampling estimate degradation. Our findings are consistent with previous studies demonstrating the benefits of shorter sampling periods. Metcalf et al., (2022) showed that employing a higher number of shorter sampling periods, rather than fewer longer sampling periods, improved species richness estimates for bird communities in tropical forests. Francomano et al., (2021) also found that using a larger number of equally spaced subsampling divisions within a fixed time period reduced subsampling error when estimating three acoustic indices of biodiversity, compared to using fewer longer subsampling divisions. Our findings also align with PAM studies in marine environments, which found that short cycle lengths in duty-cycle subsampling provided the most reliable estimates of daily acoustic presence and call rates, for North Atlantic right whale and Antarctic blue whale (Thomisch et al., 2015), as well as Southern Resident killer whale populations (Rand et al., 2022). Shorter cycle lengths additionally offer better temporal resolution in describing fluctuations of the monitored quantity. However, if the cycle length is too short, estimates of the monitored quantity may become noisy (Shimazaki and Shinomoto, 2007), and more transient biological activities, such as bat feeding buzzes, may not be captured adequately. Battery consumption would also increase as a result of switching between ON/OFF states more frequently when cycle lengths are shortened.

Comparing results across a variety of duty-cycle subsampling schemes allowed us to isolate the associated effects into cycle length and listening ratio. The optimal combination of cycle length and listening ratio ultimately depends on the exact monitoring scenario, including the behaviors and abundance of the target species or taxa, desired temporal resolution, and practical equipment and logistical constraints.

Our simulation further showed that duty-cycle subsampling estimates are more accurate when the overall call activity is high (Fig. 6C, Fig. S14, Fig. S15). This can be explained by the greater probability of detecting calls during the recorder ON period in a cycle, as bat calls occur in bouts spanning durations on the order of seconds, which can be easily missed if bouts are rare. When the overall call activity is high, metrics estimated using calls detected during the recorder ON period are also likely more representative of the true values, given the well-known challenge to accurately quantify rare events (Shimazaki and Shinomoto, 2007). Our results are consistent with a previous finding that sampling intensity needed to detect bat population declines can be lowered if the initial occupancy rate (the proportion of sites where a species is expected to occur) is high (Baumgardt et al., 2022). Another study similarly found that the acoustic complexity index (ACI) can be more accurately estimated under duty-cycle subsampling when the call activity exhibits less temporal fluctuation (Pieretti et al., 2015). ACI is a metric that quantifies general species diversity by measuring the amount of variation in acoustic intensity within a frequency bin during a given time interval. In that study, ACI estimates from nighttime insect choruses, which are more constant across time, required less intensive sampling schedules than ACI estimates from daytime bird songs, which exhibit greater temporal variability.

Lastly, comparison across the phonic groups showed that LF call activity metrics are estimated more accurately than HF call activity metrics under duty-cycle subsampling (Fig. 6D, Fig. S16). Similar to the comparison across activity levels, this can be explained by the relative abundance of LF call detections compared to HF call activity. Multiple factors contribute to LF calls being more likely detected than HF calls (Kaiser and O’Keefe, 2015). First, for calls with the same source level, LF calls travel farther than HF calls due to the stronger atmospheric absorption and reduced scattering from objects in the environment at higher frequencies (Kinsler, 2000; Stilz and Schnitzler, 2012). Similar differences have been cited to argue for longer cycle lengths in the PAM of blue whales than killer whales, due to their significantly lower call frequency content (Rand et al., 2022; Thomisch et al., 2015). Second, Carp Pond LF calls occupied a larger proportion of recording cycles than Telephone Field HF calls. This resulted from the LF phonic group having longer bout durations while maintaining BBIs similar to the HF phonic group (Fig. 5B-C). Lastly, the AudioMoth microphone response is better at the frequency range of LF calls (Lapp et al., 2023).

Across the three activity metrics, AI and BTP appear to be more robust than CR under duty-cycle subsampling. Both AI and BTP are temporally aggregated metrics, while CR is a “raw” metric that is more sensitive to any changes in the number of detected calls (Fig. 3A). Within the recorder ON period, AI represents the percentage of time blocks with calls present (Fig. 3B), and BTP captures the percentage of time period spanned by closely spaced calls (Fig. 3C). While these two measures are similar, AI is discrete in nature due to the fixed length of the time blocks (5 secs in this study), whereas BTP would change according to the actual timing of call occurrence. Therefore, duty-cycle estimates of AI are likely more invariant than duty-cycle estimates of BTP despite AI’s lower temporal resolution, as shown in the simulation results (Fig. 6, Fig. S12, Fig. S13). Similarly, duty-cycle estimates of AI outperformed duty-cycle estimates of BTP for HF calls than LF calls (Fig. 6D, Fig. S16), due to the higher temporal variability of HF calls (shorter bouts, Fig. 5B).

When false positives are present, estimates of the three metrics behave differently depending on the overall call activity (Fig. 7A-C, solid vs dashed lines). When the overall call activity is high, AI is relatively insensitive to false positives, because many time blocks would have already been occupied with the original calls. When the overall call activity is low, however, AI is susceptible to false positives, because some of the originally empty blocks would become occupied, elevating the resulting AI. On the other hand, BTP is in general relatively insensitive to false positives, because the bout duration would increase only when the false positives occur immediately adjacent to (with a spacing <BCI) an existing bout. When the overall activity is low and the false positives are singularly distributed across time, they do not form bouts and therefore do not affect BTP. CR is the most impacted by false positives when the call activity is high, as false positives directly alter the number of calls. However, note that when the overall call activity is low, singularly distributed false positives do not increase CR estimates as substantially as AI estimates.

Under the impact of false negatives, estimates of the three metrics also behave differently depending on the overall call activity (Fig. 7D-F). When the overall call activity is high, AI is insensitive to false negatives (missed calls), which likely occur within time blocks already occupied by many other calls. Similarly, BTP changes would also be small, because the bout duration only changes slightly, on the order of milliseconds to seconds, when the false negatives occur at the edges of existing bouts, or when they break an existing bout into two. These cases become more likely when the overall call activity is low. When the overall call activity is low, AI is more susceptible to false negatives, because some time blocks may be occupied by only very few calls that are falsely missed, resulting in a lower AI. Similarly, BTP is more susceptible to false negatives, as bouts are less likely to be formed when many calls are missed. Similar to the scenarios discussed for false positives (the previous paragraph), CR is the most impacted by false negatives when the call activity is high, as false negatives directly alter the number of detected calls. However, note that all three metrics perform equally poorly when the overall call activity is low.

The results we present in this study may be affected by the bat call detectors we used and the content of the dataset itself. All results included in the main text were based on simulated data generated by treating calls detected `batdetect2` as “true” call activity contained in our data. `batdetect2`, like all automated detectors, does produce false positives and missed calls (see Supplemental Materials Section S1). However, additional analyses using Kaleidoscope Pro call detections produced the same patterns in terms of the quality of duty-cycle estimates (Fig. S4, Fig. S5), despite the very different error characteristics of these two detectors (Table S1), suggesting the reliability of our results. In addition, our dataset was collected in a temperate ecosystem with a much simpler species composition than tropical regions. Greater species diversity in tropical regions likely results in a wider range of acoustic events and more varied calls, potentially affecting detector error characteristics and, consequently, estimates of activity metrics. Differences in temperature and humidity may also influence the call detection range and in turn impact the call detection probability. Future studies using datasets from tropical regions would shed light on how these differences influence duty-cycle subsampling in bat PAM.

In addition, in this study we successfully applied the concept of a “bout” (Slater and Lester, 1982) to automatically calculate the time spanned by a sequence of closely spaced bat calls (see Supplemental Materials Section S3). This sequence is similar to a “bat pass,” which has been defined in the literature as a sequence of 2 or more echolocation calls recorded as a single bat flies through air space (Fenton, 1970). Bat passes have been used to quantify activity level, species-specific temporal call activity pattern, and time spent foraging within the detection volume of the recorder (Beason et al., 2020; Brodgers, 2003; Froidevaux et al., 2023; Kerbiriou et al., 2019; Kunz et al., 2007; Rautenbach et al., 1996; Tuneu-Corral et al., 2020), and have also been identified by fixed recording intervals of 5-s containing bat calls (Millon et al., 2015) or through manual annotation by experts, which may introduce subjective bias in the grouping (Miller, 2001). In this study, we use bouts to capture sequences of bat calls detected in a recording, similar to the duration of bat passes defined in Kerbiriou et al., (2019). By associating bouts with bat passes, here we provide a method to objectively and automatically identify bat passes in long-term bat PAM.

## V. CONCLUSION

In this study, we systematically investigated the effects of duty-cycle subsampling configurations under different monitoring conditions and the performance of three activity metrics given different subsampling and detector error characteristics. We found that shorter cycle lengths and larger listening ratios are better under duty-cycle subsampling, and that subsampled data provide better activity estimates when the probability of detecting calls is high—either when calls are abundant or when call frequencies are lower. In addition, we found that the two temporally aggregated activity metrics (AI and BTP) are in general more robust to subsampling and detector errors than the absolute metric (CR). Our analysis approach, which employed simulated data by subsampling continuously recorded data, further provides a framework that can be adapted to explore other subsampling schemes in different monitoring environments.

We summarize our findings as a set of reference guidelines (Table 1) for selecting duty-cycle subsampling and activity metrics for future bat PAM design. Specifically, our results for the effects of cycle length and listening ratio provide a basis for designing subsampling schemes for different monitoring conditions. For example, where activity level and frequency of detectable calls fluctuate over a long time period, subsampling schemes can be adjusted to conserve battery and storage capacity, as recommended in Baumgardt et al., (2022). Our results on the performance of CR, AI, and BTP underscore the importance of selecting activity metrics that best suit the nature of the research questions and are robust to the error characteristics of automated detectors. For example, whether temporally aggregated metrics (such as AI and BTP) or absolute measures (such as CR) are more suitable in detecting trends in habitat usage (Biffi et al., 2024; Dietzer et al., 2024) or population declines such as due to white-nose syndrome (WNS) (Hicks et al., 2020), and whether the detectors are more prone to produce false detections or missing calls, as these factors may impact downstream analyses and interpretation (Clement et al., 2022; Miller et al., 2011; Singer et al., 2024). The results and approach presented here can help support PAM programs worldwide by balancing information retention, error tolerance, and logistical resources, such as equipment availability and costs.

**TABLE I.**
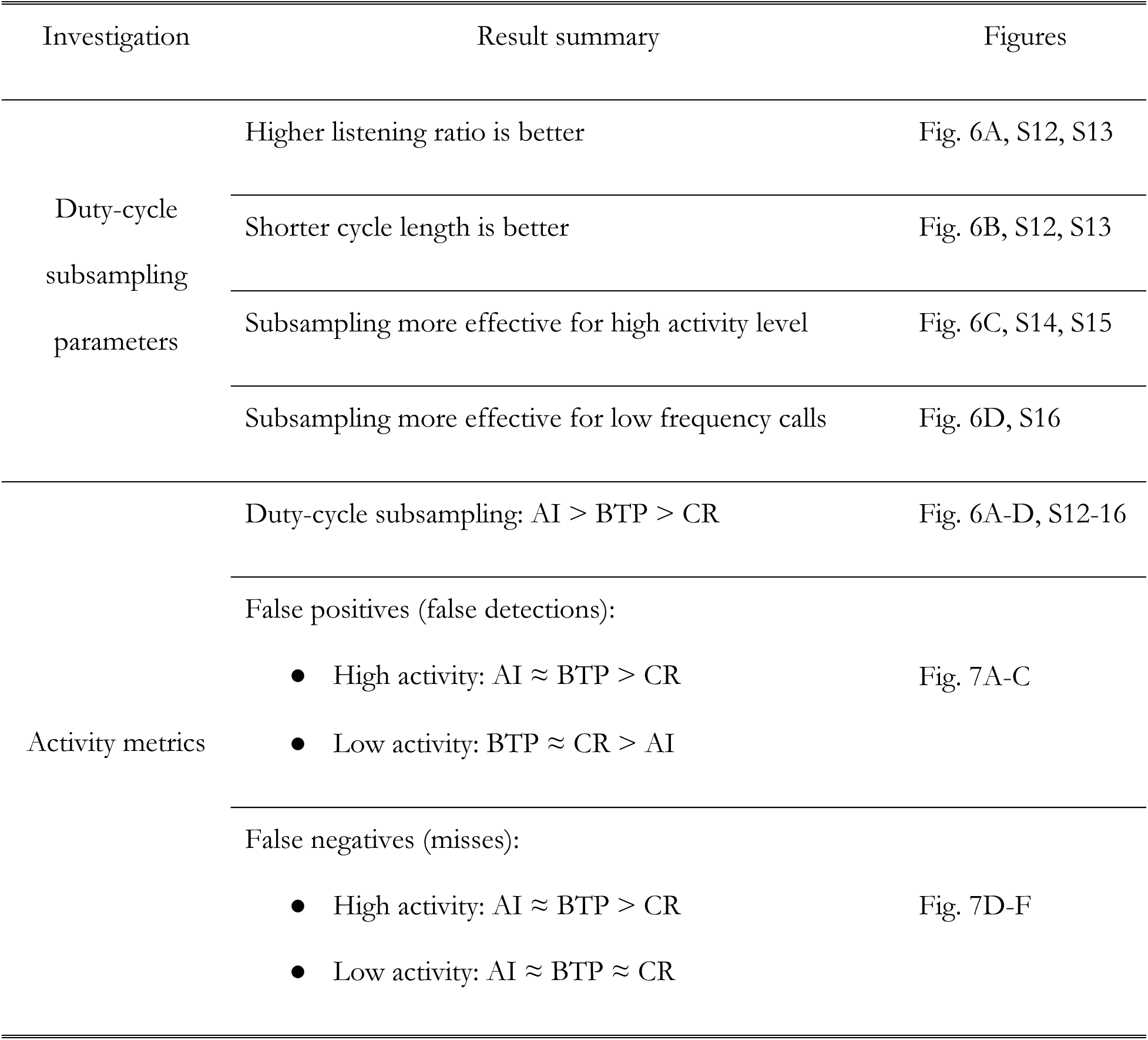
Guidelines for selecting duty-cycle subsampling configuration and activity metrics in bat PAM based on findings in this study.

## SUPPLEMENTARY MATERIAL

See supplementary material at [URL will be inserted by AIP] for supplementary text, figures, and tables.

## ACKNOWLEDGMENTS

We thank Josie Sachen and YeonJoon Cheong for assisting with data collection, and Corbin Charpentier, Kirsteen Ng, and Ernesto Cediel for developing the software pipeline as part of a Master’s in Data Science Capstone project. We thank funding support from the University of Washington (UW) Mary Gates Endowment for Students, UW Royalty Research Funds, UW Applied Physics Laboratory, and UW Department of Electrical and Computer Engineering, and the Office of Naval Research (ONR) Multidisciplinary University Research Initiatives (MURI) Program grant N00014-23-12065. We also thank the Open Storage Network (OSN) for providing cloud data storage services under allocation BIO230143 from the Advanced Cyberinfrastructure Coordination Ecosystem: Services & Support (ACCESS) program, which is supported by National Science Foundation grants #2138259, #2138286, #2138307, #2137603, and #2138296.

## AUTHOR CONTRIBUTIONS

W.-J.L. conceived of the idea and worked with A.K. to design methodology. A.K. collected and analyzed the data. A.K. and W.-J.L. interpreted the results and wrote the manuscript.

## DATA AVAILABILITY

Field records during data collection have been uploaded to a publicly accessible repository: https://github.com/uw-echospace/ubna-field

Code for data processing and analyses to generate all paper results are also hosted in a publicly accessible repository: https://github.com/uw-echospace/duty-cycle-analysis

The AudioMoth recordings that support the findings of this study are available from the corresponding author upon reasonable request due to the very large data volume.

## REFERENCES

Anunciação, P. R., Sugai, L. S. M., Martello, F., De Carvalho, L. M. T., and Ribeiro, M. C. (2022). “Estimating the diversity of tropical anurans in fragmented landscapes with acoustic monitoring: lessons from a sampling sufficiency perspective,” Biodivers. Conserv., 31, 3055–3074. doi:10.1007/s10531-022-02475-w

Aodha, O. M., Martínez Balvanera, S., Damstra, E., Cooke, M., Eichinski, P., Browning, E., Barataud, M., et al. (2022). “Towards a General Approach for Bat Echolocation Detection and Classification.,” doi:10.1101/2022.12.14.520490

Atkin, P (2024). “π•pistrelle Bat Detector” https://www.pippyg.com (Last viewed April 11, 2025)

Baumgardt, J. A., Morrison, M. L., Brennan, L. A., Davis, H. T., Fern, R. R., Szewczak, J. M., and Campbell, T. A. (2022). “Monitoring Occupancy of Bats with Acoustic Data: Power and Sample Size Recommendations,” West. North Am. Nat., , doi: 10.3398/064.082.0104. doi:10.3398/064.082.0104

Beason, R. D., Riesch, R., and Koricheva, J. (2020). “Temporal Pass Plots: An intuitive method for visualising activity patterns of bats and other vocalising animals,” Ecol. Indic., 113, 106202. doi:10.1016/j.ecolind.2020.106202

Biffi, S., Chapman, P. J., Engler, J. O., Kunin, W. E., and Ziv, G. (2024). “Using automated passive acoustic monitoring to measure changes in bird and bat vocal activity around hedgerows of different ages,” Biol. Conserv., 296, 110722. doi:10.1016/j.biocon.2024.110722

Brinkløv, S. M. M., Macaulay, J., Bergler, C., Tougaard, J., Beedholm, K., Elmeros, M., and Madsen, P. T. (2023). “Open-source workflow approaches to passive acoustic monitoring of bats,” Methods Ecol. Evol., 14, 1747–1763. doi:10.1111/2041-210X.14131

Brodgers, H. G. (2003). “Another Quantitative Measure of Bat Species Activity and Sampling Intensity Considerations for the Design of Ultrasonic Monitoring Studies,” Acta Chiropterologica, 5, 235. doi:10.3161/001.005.0206

Clement, M. J., Royle, J. A., and Mixan, R. J. (2022). “Estimating occupancy from autonomous recording unit data in the presence of misclassifications and detection heterogeneity,” Methods Ecol. Evol., 13, 1719–1729. doi:10.1111/2041-210X.13895

Cook, A., and Hartley, S. (2018). “Efficient sampling of avian acoustic recordings: intermittent subsamples improve estimates of single species prevalence and total species richness,” Avian Conserv. Ecol., 13, art21. doi:10.5751/ACE-01221-130121

Dietzer, M. T., Keicher, L., Kohles, J. E., Hurme, E. R., Ruczyński, I., Borowik, T., Zegarek, M., et al. (2024). “High temporal resolution data reveal low bat and insect activity over managed meadows in central Europe,” Sci. Rep., 14, 7498. doi:10.1038/s41598-024-57915-0

Fagen, R. M., and Young., D. Y. (1978). “Temporal patterns of behavior: durations, intervals, latencies and sequences,” In P. W. Colgan (Ed.), Quant. Ethol., Wiley, New York, pp. 79–114.

Fenton, M. B. (1970). “A technique for monitoring bat activity with results obtained from different environments in southern Ontario,” Can. J. Zool., 48, 847–851. doi:10.1139/z70-148

Francomano, D., Gottesman, B. L., and Pijanowski, B. C. (2021). “Biogeographical and analytical implications of temporal variability in geographically diverse soundscapes,” Ecol. Indic., 121, 106794. doi:10.1016/j.ecolind.2020.106794

Fraser, E. E., Silvis, A., Brigham, R. M., and Czenze, Z. J. (2020). Bat Echolocation Research - A handbook for planning and conducting acoustic studies,.

Froidevaux, J. S. P., Jones, G., Kerbiriou, C., and Park, K. J. (2023). “Acoustic activity of bats at power lines correlates with relative humidity: a potential role for corona discharges,” Proc. R. Soc. B Biol. Sci., 290, 20222510. doi:10.1098/rspb.2022.2510

Froidevaux, J. S. P., Zellweger, F., Bollmann, K., and Obrist, M. K. (2014). “Optimizing passive acoustic sampling of bats in forests,” Ecol. Evol., 4, 4690–4700. doi:10.1002/ece3.1296

Gibb, R., Browning, E., Glover-Kapfer, P., and Jones, K. E. (2019). “Emerging opportunities and challenges for passive acoustics in ecological assessment and monitoring,” (L. Börger, Ed.) Methods Ecol. Evol., 10, 169–185. doi:10.1111/2041-210X.13101

Hauer, C. L., Shinskie, J. L., Brady, R. J., and Titus, C. N. (2023). “Sampling Duration and Season Recommendations for Passive Acoustic Monitoring of Bats after White-Nose Syndrome,” J. Fish Wildl. Manag., 14, 365–384. doi:10.3996/JFWM-23-021

Hicks, L. L., Schwab, N. A., Homyack, J. A., Jones, J. E., Maxell, B. A., and Burkholder, B. O. (2020). “A statistical approach to white-nose syndrome surveillance monitoring using acoustic data,” (K. Root, Ed.) PLOS ONE, 15, e0241052. doi:10.1371/journal.pone.0241052

Hill, A. P., Prince, P., Piña Covarrubias, E., Doncaster, C. P., Snaddon, J. L., and Rogers, A. (2018). “AudioMoth: Evaluation of a smart open acoustic device for monitoring biodiversity and the environment,” (N. Isaac, Ed.) Methods Ecol. Evol., 9, 1199–1211. doi:10.1111/2041-210X.12955

Hoggatt, M. L., Starbuck, C. A., and O’Keefe, J. M. (2024). “Acoustic monitoring yields informative bat population density estimates,” Ecol. Evol., 14, e11051. doi:10.1002/ece3.11051

Kaiser, Z. D. E., and O’Keefe, J. M. (2015). “Data acquisition varies by bat phonic group for 2 types of bat detectors when weatherproofed and paired in field settings,” Wildl. Soc. Bull., 39, 635–644. doi:10.1002/wsb.572

Kerbiriou, C., Bas, Y., Le Viol, I., Lorrilliere, R., Mougnot, J., and Julien, J. F. (2019). “Potential of bat pass duration measures for studies of bat activity,” Bioacoustics, 28, 177–192. doi:10.1080/09524622.2017.1423517

Kinsler, L. E. (Ed.) (2000). Fundamentals of acoustics, Wiley, New York, 4th ed., 1 pages.

Kloepper, L. N., Linnenschmidt, M., Blowers, Z., Branstetter, B., Ralston, J., and Simmons, J. A. (2016). “Estimating colony sizes of emerging bats using acoustic recordings,” R. Soc. Open Sci., 3, 160022. doi:10.1098/rsos.160022

Kunz, T. H., Arnett, E. B., Cooper, B. M., Erickson, W. P., Larkin, R. P., Mabee, T., Morrison, M. L., et al. (2007). “Assessing Impacts of Wind-Energy Development on Nocturnally Active Birds and Bats: A Guidance Document,” J. Wildl. Manag., 71, 2449–2486. doi:10.2193/2007-270

Lahoz-Monfort, J. J., and Magrath, M. J. L. (2021). “A Comprehensive Overview of Technologies for Species and Habitat Monitoring and Conservation,” BioScience, 71, 1038–1062. doi:10.1093/biosci/biab073

Lapp, S., Stahlman, N., and Kitzes, J. (2023). “A Quantitative Evaluation of the Performance of the Low-Cost AudioMoth Acoustic Recording Unit,” Sensors, 23, 5254. doi:10.3390/s23115254

López-Baucells, A., Yoh, N., Rocha, R., Bobrowiec, P. E. D., Palmeirim, J. M., and Meyer, C. F. J. (2021). “Optimizing bat bioacoustic surveys in human-modified Neotropical landscapes,” Ecol. Appl., 31, e02366. doi:10.1002/eap.2366

Melo, I., Llusia, D., Bastos, R. P., and Signorelli, L. (2021). “Active or passive acoustic monitoring? Assessing methods to track anuran communities in tropical savanna wetlands,” Ecol. Indic., 132, 108305. doi:10.1016/j.ecolind.2021.108305

Metcalf, O. C., Barlow, J., Marsden, S., Gomes De Moura, N., Berenguer, E., Ferreira, J., and Lees, A. C. (2022). “Optimizing tropical forest bird surveys using passive acoustic monitoring and high temporal resolution sampling,” (N. Pettorelli and C. Astaras, Eds.) Remote Sens. Ecol. Conserv., 8, 45–56. doi:10.1002/rse2.227

Milchram, M., Suarez-Rubio, M., Schröder, A., and Bruckner, A. (2020). “Estimating population density of insectivorous bats based on stationary acoustic detectors: A case study,” Ecol. Evol., 10, 1135–1144. doi:10.1002/ece3.5928

Miller, B. (2001). “A method for determining relative activity of free flying bats using a new activity for acoustic monitoring,” Acta Chiropterologica, 3, 93–105.

Miller, D. A., Nichols, J. D., McClintock, B. T., Grant, E. H. C., Bailey, L. L., and Weir, L. A. (2011). “Improving occupancy estimation when two types of observational error occur: non-detection and species misidentification,” Ecology, 92, 1422–1428. doi:10.1890/10-1396.1

Millon, L., Julien, J.-F., Julliard, R., and Kerbiriou, C. (2015). “Bat activity in intensively farmed landscapes with wind turbines and offset measures,” Ecological Engineering, 75, 250–257. doi:10.1016/j.ecoleng.2014.11.050

Moss, C. F., Ortiz, S. T., and Wahlberg, M. (2023). “Adaptive echolocation behavior of bats and toothed whales in dynamic soundscapes,” J. Exp. Biol., 226, jeb245450. doi:10.1242/jeb.245450

Ober, H. K., and Hayes, J. P. (2008). “Influence of Vegetation on Bat Use of Riparian Areas at Multiple Spatial Scales,” J. Wildl. Manag., 72, 396–404. doi:10.2193/2007-193

Pieretti, N., Duarte, M. H. L., Sousa-Lima, R. S., Rodrigues, M., Young, R. J., and Farina, A. (2015). “Determining Temporal Sampling Schemes for Passive Acoustic Studies in Different Tropical Ecosystems,” Trop. Conserv. Sci., 8, 215–234. doi:10.1177/194008291500800117

Rand, Z. R., Wood, J. D., and Oswald, J. N. (2022). “Effects of duty cycles on passive acoustic monitoring of southern resident killer whale (Orcinus orca) occurrence and behavior,” J. Acoust. Soc. Am., 151, 1651–1660. doi:10.1121/10.0009752

Ratcliffe, J. M., Elemans, C. P. H., Jakobsen, L., and Surlykke, A. (2013). “How the bat got its buzz,” Biol. Lett., 9, 20121031. doi:10.1098/rsbl.2012.1031

Rautenbach, I. L., Whiting, M. J., and Fenton, M. B. (1996). “Bats in riverine forests and woodlands: a latitudinal transect in southern Africa,” Can. J. Zool., 74, 312–322. doi:10.1139/z96-039

Riera, A., Ford, J. K., and Ross Chapman, N. (2013). “Effects of different analysis techniques and recording duty cycles on passive acoustic monitoring of killer whales,” J. Acoust. Soc. Am., 134, 2393–2404. doi:10.1121/1.4816552

Ross, S. R. P. -J., O’Connell, D. P., Deichmann, J. L., Desjonquères, C., Gasc, A., Phillips, J. N., Sethi, S. S., et al. (2023). “Passive acoustic monitoring provides a fresh perspective on fundamental ecological questions,” Funct. Ecol., 37, 959–975. doi:10.1111/1365-2435.14275

Russo, D., Ancillotto, L., and Jones, G. (2018). “Bats are still not birds in the digital era: echolocation call variation and why it matters for bat species identification,” Can. J. Zool., 96, 63–78. doi:10.1139/cjz-2017-0089

Schowalter, D. B. (1980). “Swarming, Reproduction, and Early Hibernation of Myotis lucifugus and M. volans in Alberta, Canada,” J. Mammal., 61, 350–354. doi:10.2307/1380065

Shimazaki, H., and Shinomoto, S. (2007). “A Method for Selecting the Bin Size of a Time Histogram,” Neural Comput., 19, 1503–1527. doi:10.1162/neco.2007.19.6.1503

Sibly, R. M., Nott, H. M. R., and Fletcher, D. J. (1990). “Splitting behaviour into bouts,” Anim. Behav., 39, 63–69. doi:10.1016/S0003-3472(05)80726-2

Singer, D., Hagge, J., Kamp, J., Hondong, H., and Schuldt, A. (2024). “Aggregated time-series features boost species-specific differentiation of true and false positives in passive acoustic monitoring of bird assemblages,” Remote Sens. Ecol. Conserv., 10, 517–530. doi:10.1002/rse2.385

Skalak, S. L., Sherwin, R. E., and Brigham, R. M. (2012). “Sampling period, size and duration influence measures of bat species richness from acoustic surveys: Effective acoustic monitoring,” Methods Ecol. Evol., 3, 490–502. doi:10.1111/j.2041-210X.2011.00177.x

Slater, P. J. B., and Lester, N. P. (1982). “Minimising Errors in Splitting Behaviour Into Bouts,” Behaviour, 79, 153–161. doi:10.1163/156853982X00229

Speakman, J. R., and Rowland, A. (1999). “Preparing for inactivity: How insectivorous bats deposit a fat store for hibernation,” Proc. Nutr. Soc., 58, 123–131. doi:10.1079/PNS19990017

Stilz, W.-P., and Schnitzler, H.-U. (2012). “Estimation of the acoustic range of bat echolocation for extended targets,” J. Acoust. Soc. Am., 132, 1765–1775. doi:10.1121/1.4733537

Thomisch, K., Boebel, O., Zitterbart, D. P., Samaran, F., Van Parijs, S., and Van Opzeeland, I. (2015). “Effects of subsampling of passive acoustic recordings on acoustic metrics,” J. Acoust. Soc. Am., 138, 267–278. doi:10.1121/1.4922703

Tuneu-Corral, C., Puig-Montserrat, X., Flaquer, C., Mas, M., Budinski, I., and López-Baucells, A. (2020). “Ecological indices in long-term acoustic bat surveys for assessing and monitoring bats’ responses to climatic and land-cover changes,” Ecol. Indic., 110, 105849. doi:10.1016/j.ecolind.2019.105849

