## Supplemental text, figures, and tables for "Influence of duty-cycle recording on measuring bat activity in passive acoustic monitoring"

### SECTION S1: CALL DETECTION

We performed automated call detection using the `batdetect2` package, which provides users with the option to set a minimum confidence score for detection. However, in preliminary analysis we found that the detector confidence score can be variable for low SNR calls. Therefore, using 5613 human-annotated bat calls in four 30-min recording segments and a series of precision-recall curves calculated with varying SNR requirements, we empirically determined a confidence score threshold of 0.35 and an SNR threshold at 3 dB to ensure the quality of detected calls. We further observed that `batdetect2` has a tendency of detecting harmonics or different temporal segments of the same call as different calls. To remove these redundant detections, we implemented an additional filtering step to retain only the detection with the highest confidence score among all nearby detections with peak energy occurring within a 12 msec window.

We also used the human-annotated bat calls to evaluate the error characteristics of both `batdetect2` and Kaleidoscope Pro in terms of precision, recall, and F1-score. These metrics were computed using true positives (TP), false positives (FP), and false negatives (FN), by considering the human annotations as "true" calls. Precision is defined as the proportion of detections that correspond to true calls:

$$\text{Precision} = \frac{\text{TP}}{\text{TP} + \text{FP}}$$

Recall is defined as the proportion of true calls that were successfully detected:

$$\text{Recall} = \frac{\text{TP}}{\text{TP} + \text{FN}}$$

F1-score is the harmonic mean of the detector's precision and recall that summarizes the overall detector performance:

$$\text{F1-score} = \frac{2 * \text{precision} * \text{recall}}{\text{precision} + \text{recall}}$$

The results show that `batdetect2` has higher precision, recall, and F1-score than Kaleidoscope Pro (Table S1), and that Kaleidoscope Pro produced many missed calls (false negatives) and produced a decent number of false detections (false positives).

TABLE SI. Detector performance of `batdetect2` and Kaleidoscope Pro.

| Detector | Precision | Recall | F-1 Score |
|---|---|---|---|
| `batdetect2` | 0.911 | 0.796 | 0.850 |
| Kaleidoscope Pro | 0.607 | 0.13 | 0.214 |

### SECTION S2: CALL CLASSIFICATION

We observed that the *k*-means classifier misclassified a small percentage of bat calls because certain LF calls contained significant energy in the higher frequency band, despite these calls clearly being part of an LF call sequence. We removed these misclassified calls by comparing the peak frequency of each detection with the median peak frequency of all calls classified into the same group within the same 30-min recording segment. This results in a small percentage of calls dropped across all 30-min recording segments (Fig. S1). On average, files with more than 20% calls dropped consisted of less than 100 calls.

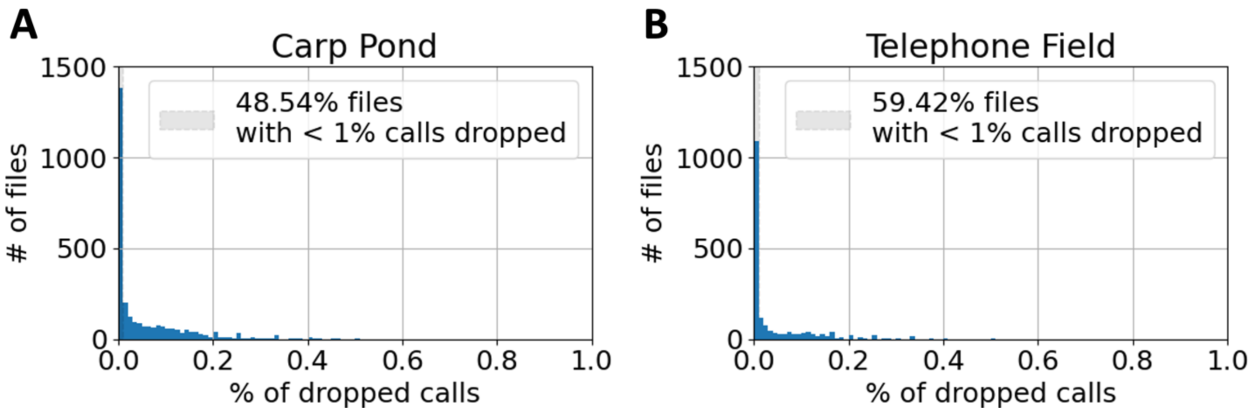


FIG. S1. Histograms showing the distribution of the number of recording segments with respect to the percentage of calls removed due to misclassification.

### SECTION S3: DETERMINATION OF CALL BOUTS

Grouping calls into bouts requires identifying the bout criterion interval (BCI), the maximum allowable time gap between any two consecutive calls for them to be grouped into the same bout. Slater and Lester (1982) proposed to identify the BCI by modeling the log-survivorship curves of the call inter-pulse intervals (IPIs) through two interval generating processes, one generating the within-bout intervals (WBIs) and the other generating the between-bout intervals (BBIs). These two processes can be modeled by performing linear regression on a segment of the log-survivorship curve representing candidates of each process, and the intersection of these processes is the BCI (Fig. S2).

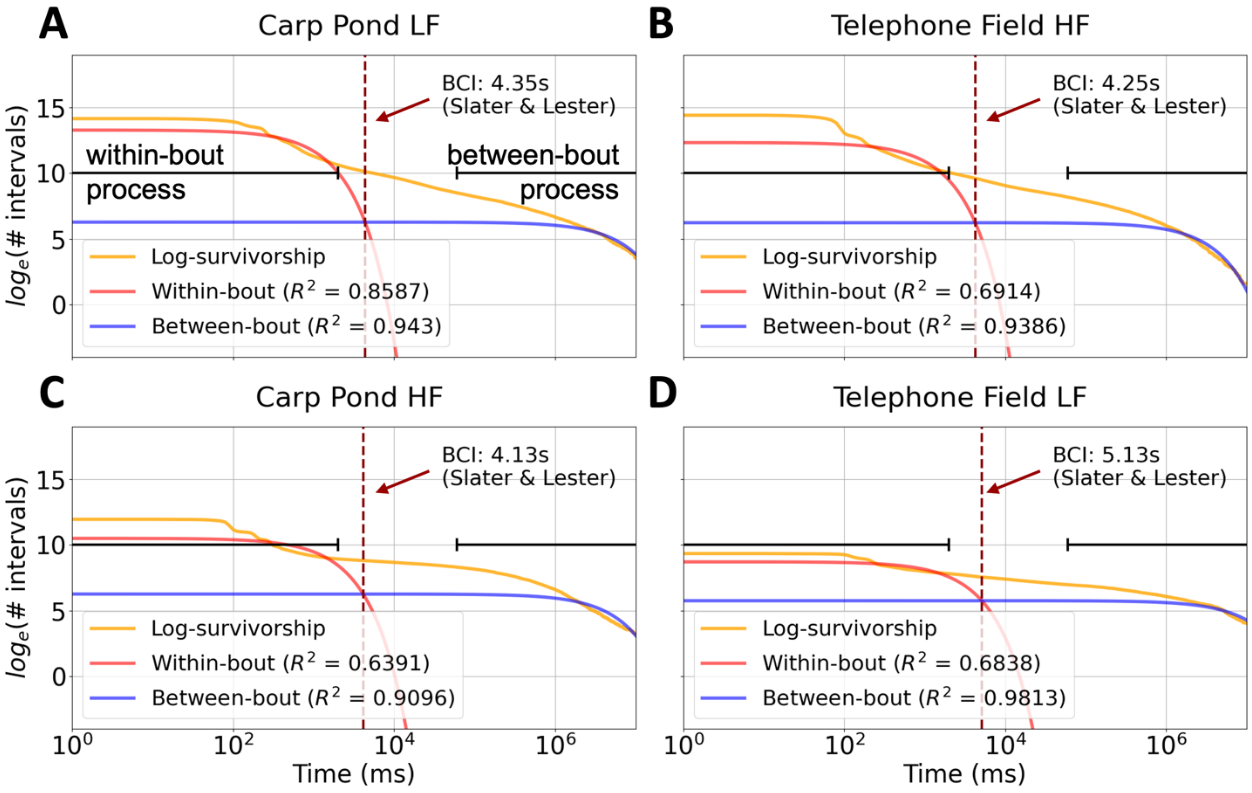


FIG. S2. Log-survivorship curves of the call inter-pulse intervals (IPIs) for the LF and HF phonic groups at Carp Pond and Telephone Field.

### SECTION S4: UPPER BOUNDS OF DUTY-CYCLE SUBSAMPLING ESTIMATES

The duty-cycle subsampling estimates of call rate (CR), activity index (AI), and bout-time percentage (BTP) each has an upper limit in terms of its deviation from the true values (Fig. 4, dashed lines). This upper limit arises from the assumption that call activity observed during the recorder ON time is representative of call activity during the entire cycle length ($\mathrm{CL}$). This assumption is most incorrect when all call activity for a given $\mathrm{CL}$ occurs only within the recorder ON period (Fig. S3). In such cases, the duty-cycle estimate becomes a multiple of the true value, scaled by the listening ratio ($\mathrm{LR}$). An example of this can be seen in Fig. S3, where call activity occurs within the first 10 minutes of a 30-min recording segment. Assuming this call activity consists of $C$ calls, occupies $A$ number of 5-sec blocks, and has $B$ seconds of bouts, we can compute CR, BTP, and AI for both the continuous and duty-cycle schemes with a CL of 30 mins as shown in Table S2. This simple exercise shows that regardless of the metric used, if the true value of the metric is $M$, where $M = C, A$, or $B$, the duty-cycle estimate will be $(\mathrm{LR})^{-1} \times M$ because the amount of absolute activity ($C$, $A$, or $B$) remains the same for either the continuous and the subsampled time period.

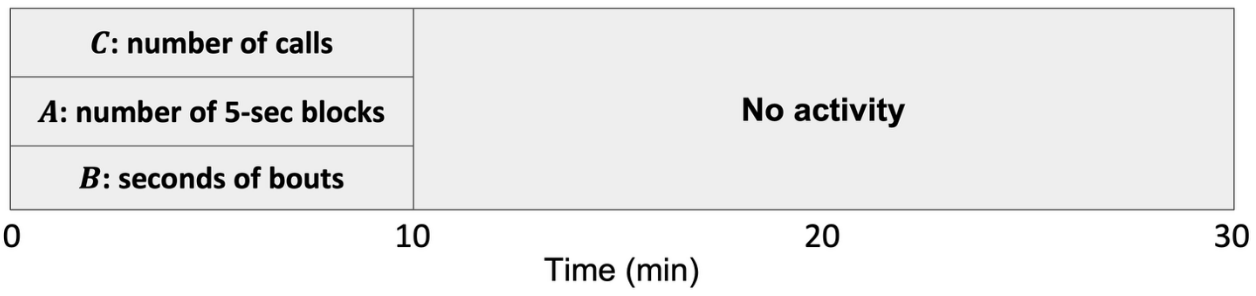


FIG. S3. Diagram showing an example time period in which call activity would be overestimated at the upper bound under duty-cycle subsampling.

TABLE S2. CR, AI, and BTP values for the example shown in Fig. S3, in which all activity occurs in the first 10 minutes of a 30-min recording segment. The first column shows the true metric values, while subsequent columns show the metric values under duty-cycle subsampling with a cycle length (CL) of 30 min and a listening ratio (LR) as shown in the column heading. $M$: true value of the metric ($M = A, B$, or $C$).

| $\mathrm{CL} = 30$ | True value | LR = $2/3$ | LR = $1/2$ | LR = $1/3$ |
|---|---|---|---|---|
| CR (calls/min) | $C/30$ | $C/20$ | $C/15$ | $C/10$ |
| AI (%) | $A/360$ | $A/240$ | $A/180$ | $A/120$ |
| BTP (%) | $B/1800$ | $B/1200$ | $B/900$ | $B/600$ |
| **Metric** | $M$ | $\frac{3}{2}M$ | $2M$ | $3M$ |

### SECTION S5: DUTY-CYCLE SUBSAMPLING SIMULATION USING KALEIDOSCOPE PRO DETECTIONS

To verify that our results are not idiosyncratic of `batdetect2` detections, we conducted an identical set of duty-cycle subsampling analyses using calls detected using Kaleidoscope Pro, The Kaleidoscope Pro call detections were configured to search for acoustic events within the 15–120 kHz frequency range, with a detected pulse length of 0.5-100 ms, a maximum inter-syllable gap of 20 ms, and a minimum number of 1 pulse per detection. Detections were visually examined to confirm that they are individual bat search-phase calls for a subset of files.

The simulated duty-cycle subsampling results derived from Kaleidoscope Pro call detections are in general consistent with those from `batdetect2` (Fig. S3). However, detailed comparison shows an overall reduction of performance across all activity metrics for Kaleidoscope estimates (Fig. S4). This is likely due to the lower overall call activity resulting from Kaleidoscope Pro's tendency to miss calls. Recall that the quality of subsampling estimates degrades when the call activity is lower or less constant (see Sec. IV).

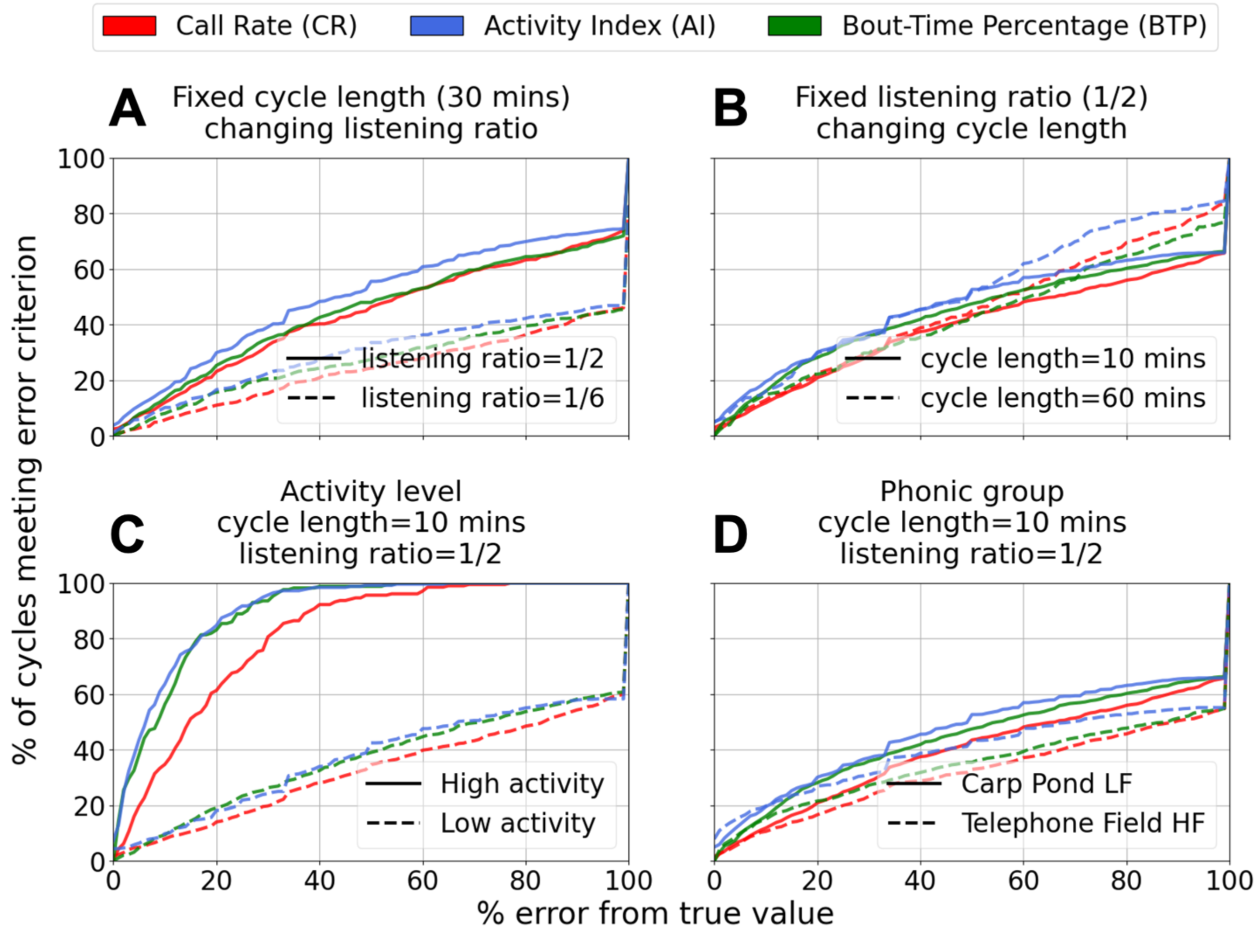


FIG. S4. Quality of duty-cycle subsampling estimates using Kaleidoscope Pro call detections. The duty-cycle parameters used here are the same as those used in Fig. 6.

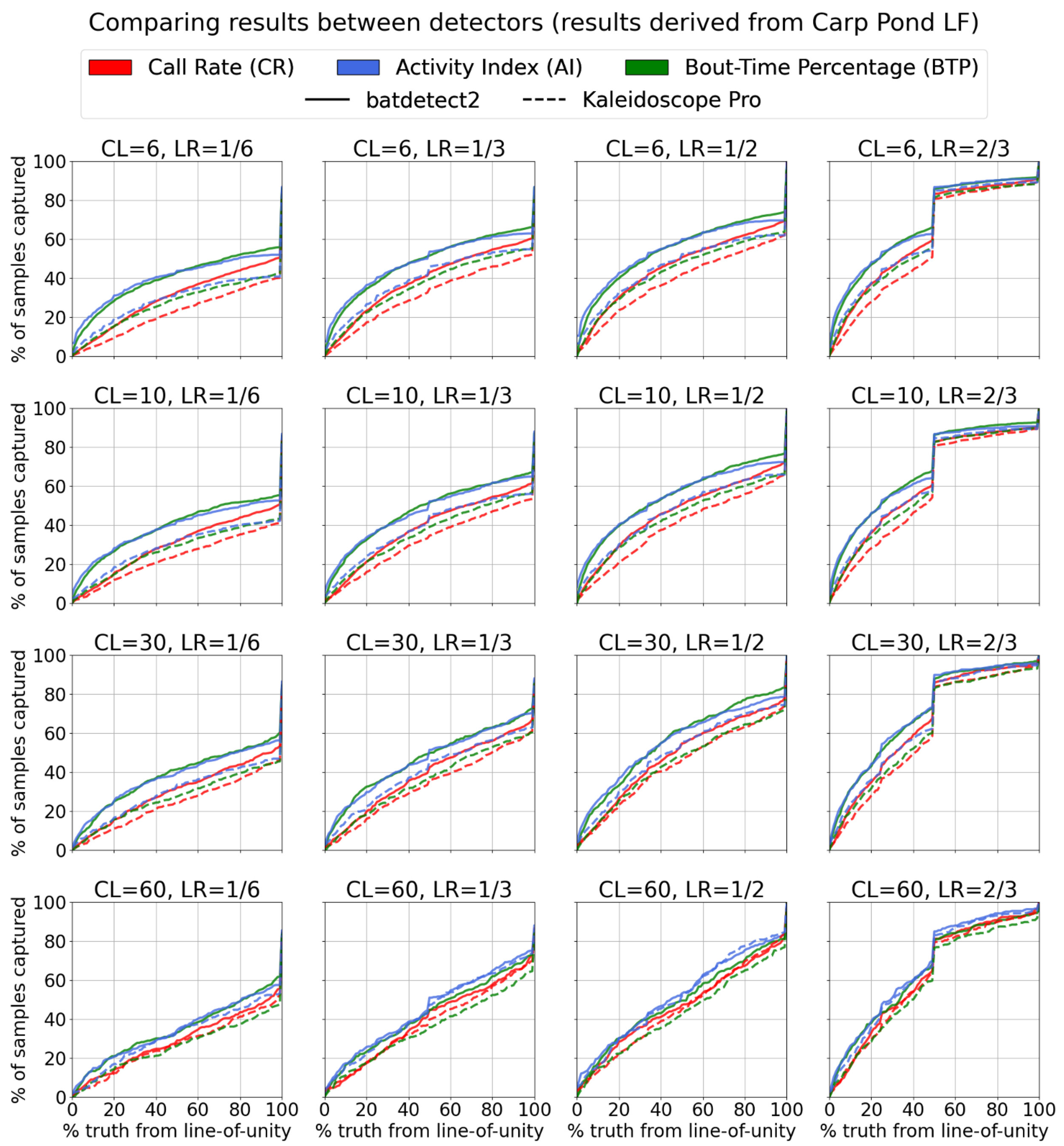


FIG. S5. Comparison of the influence of duty-cycle subsampling using `batdetect2` and Kaleidoscope Pro detections of LF calls in Carp Pond during July-August, 2022. CL: cycle length (mins); LR: listening ratio.

### SECTION S6: ADDITIONAL SIMULATION RESULTS

As mentioned in Sec. II D., we performed duty-cycle subsampling simulations systematically across a range of CLs (6, 10, 30, 60 mins) and LRs (⅙, ⅓, ½, ⅔). Fig. 6 contains a subset of these results that demonstrate the influence of duty-cycle subsampling on estimating bat call activity across all three activity metrics. Below we include the full set of simulation results by separately presenting the distributions of duty-cycle samples for Carp Pond LF calls (Fig. S6-8) and Telephone Field HF calls (Fig. S9-11), the corresponding duty-cycle evaluation curves (Fig. S12 and Fig. S13), the influence of duty-cycle subsampling on high and low activity (Fig. S14), and the comparison between Carp Pond LF and Telephone Field HF across all investigated duty-cycle configurations (Fig. S16).

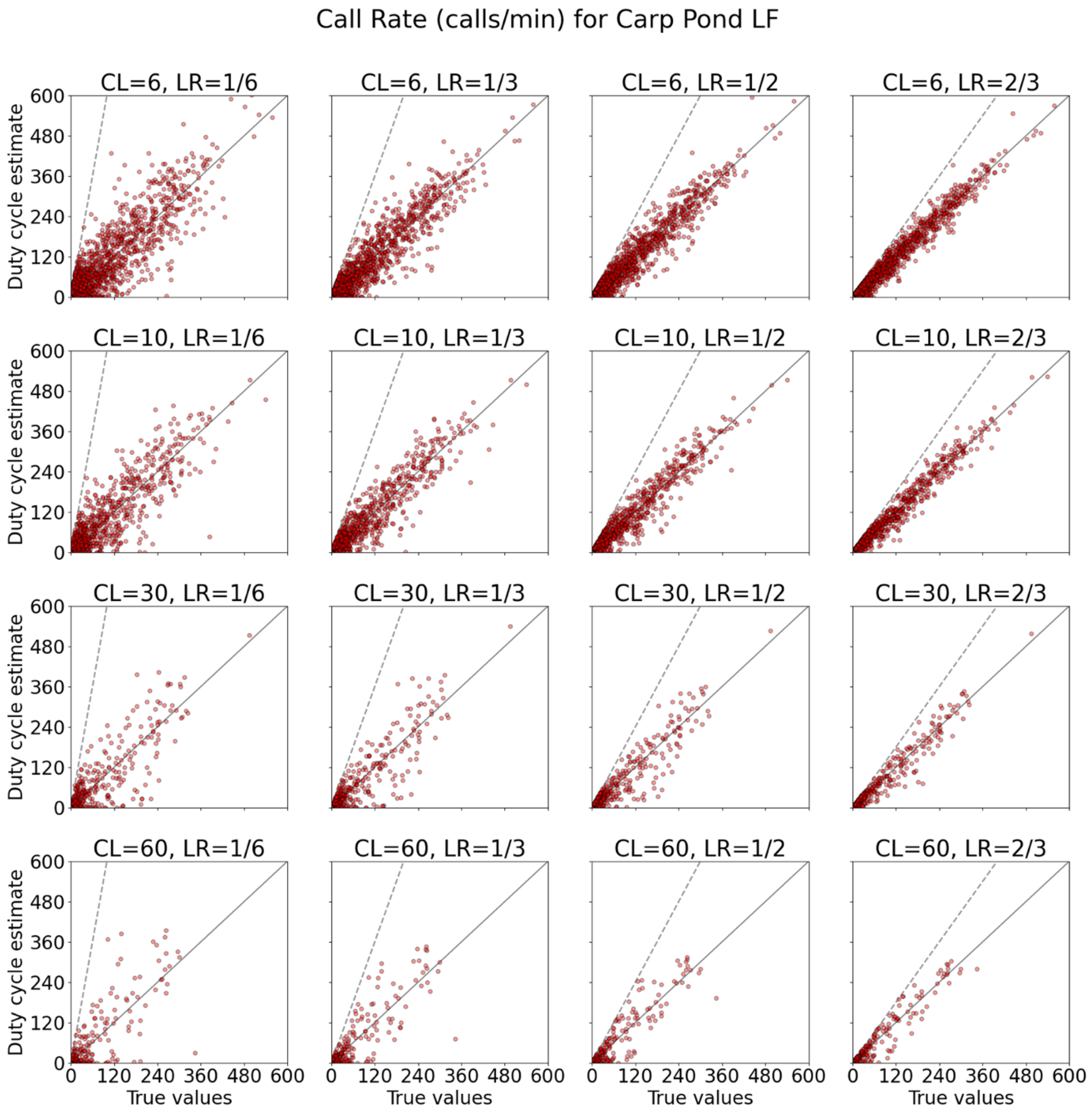


FIG. S6. Scatter plots comparing CR duty-cycle estimates with true CR values across varying duty-cycle schemes, denoted by the cycle length (CL; mins) and listening ratio (LR). The slanted grey dashed line in all panels is the upper bound of duty-cycle estimate that varies depending on the LR (see Sec. S4). The panel titled "CL=10, LR=1/2" is shown as a part of Fig. 4A. Samples closer to the line-of-unity represent when the estimate matches the true value. All panels were generated using duty-cycle samples of Carp Pond LF calls during July-August, 2022.

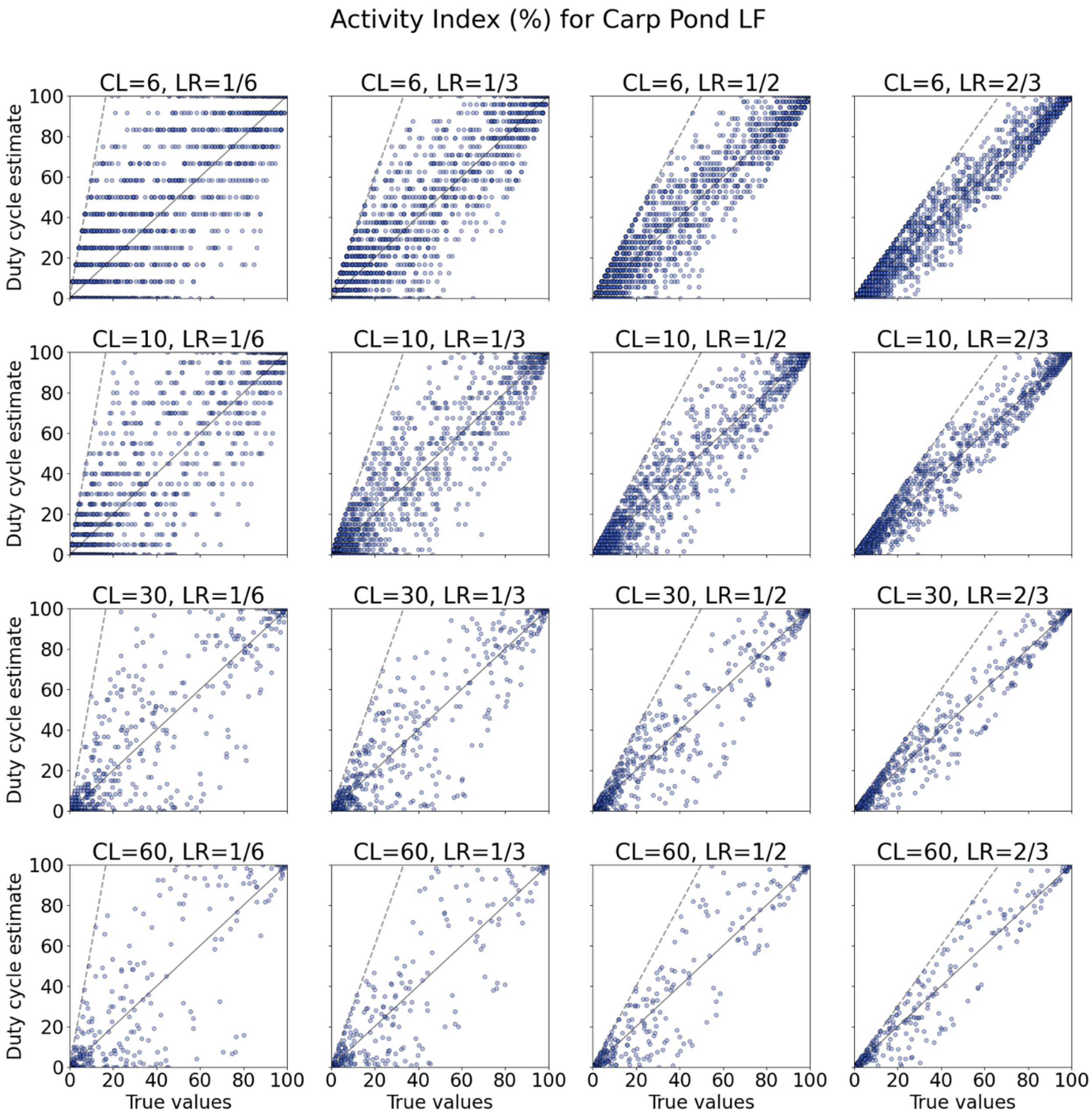


FIG. S7. Scatter plots comparing AI duty-cycle estimates with true AI values across varying duty-cycle schemes from the overall investigation. All other plot details are identical to those in Fig. S6. CL: cycle length (mins); LR: listening ratio.

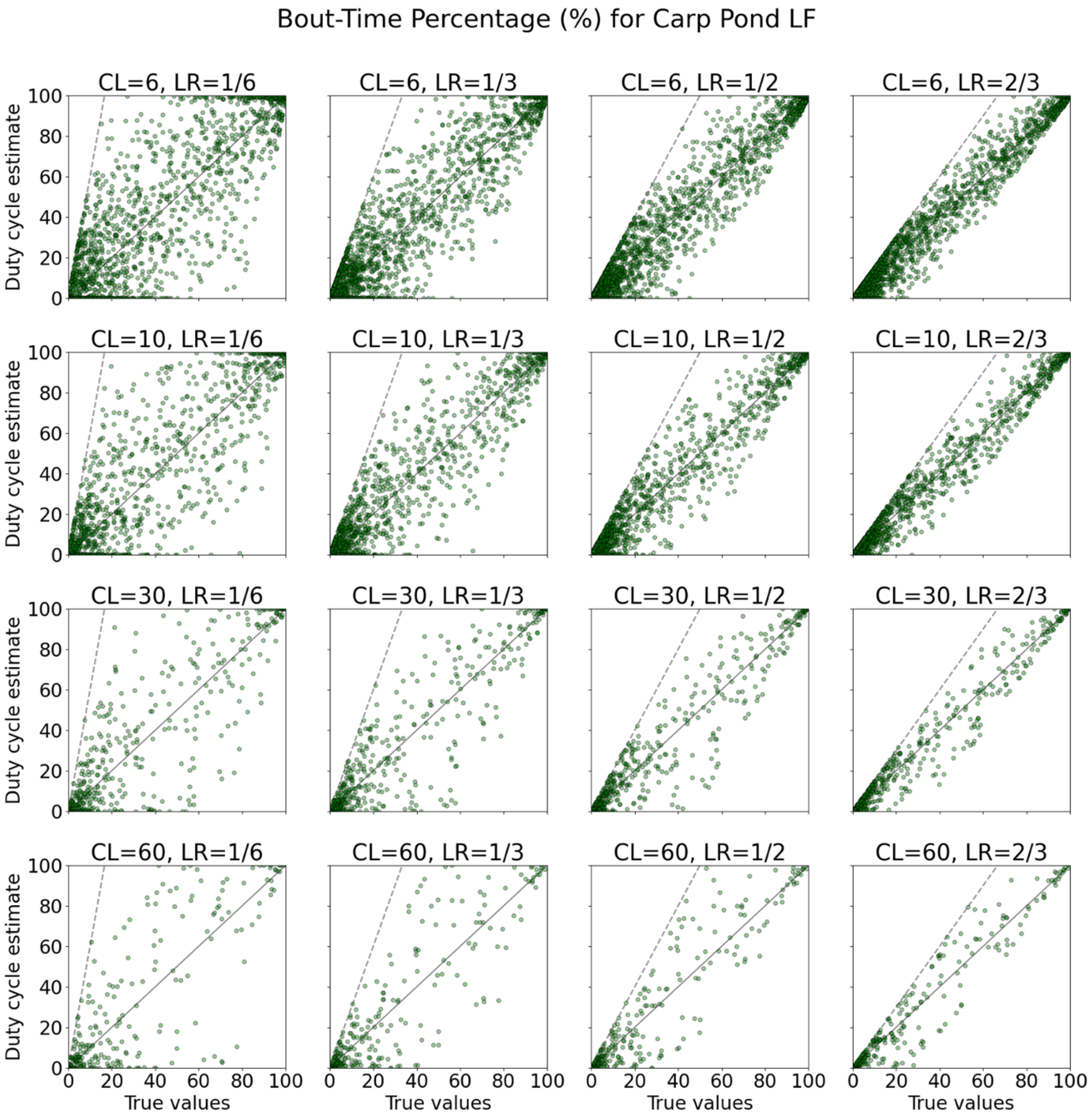


FIG. S8. Scatter plots comparing BTP duty-cycle estimates with true BTP values across varying duty-cycle schemes from the overall investigation. All other plot details are identical to those in Fig. S6. **CL**: cycle length (mins); **LR**: listening ratio.

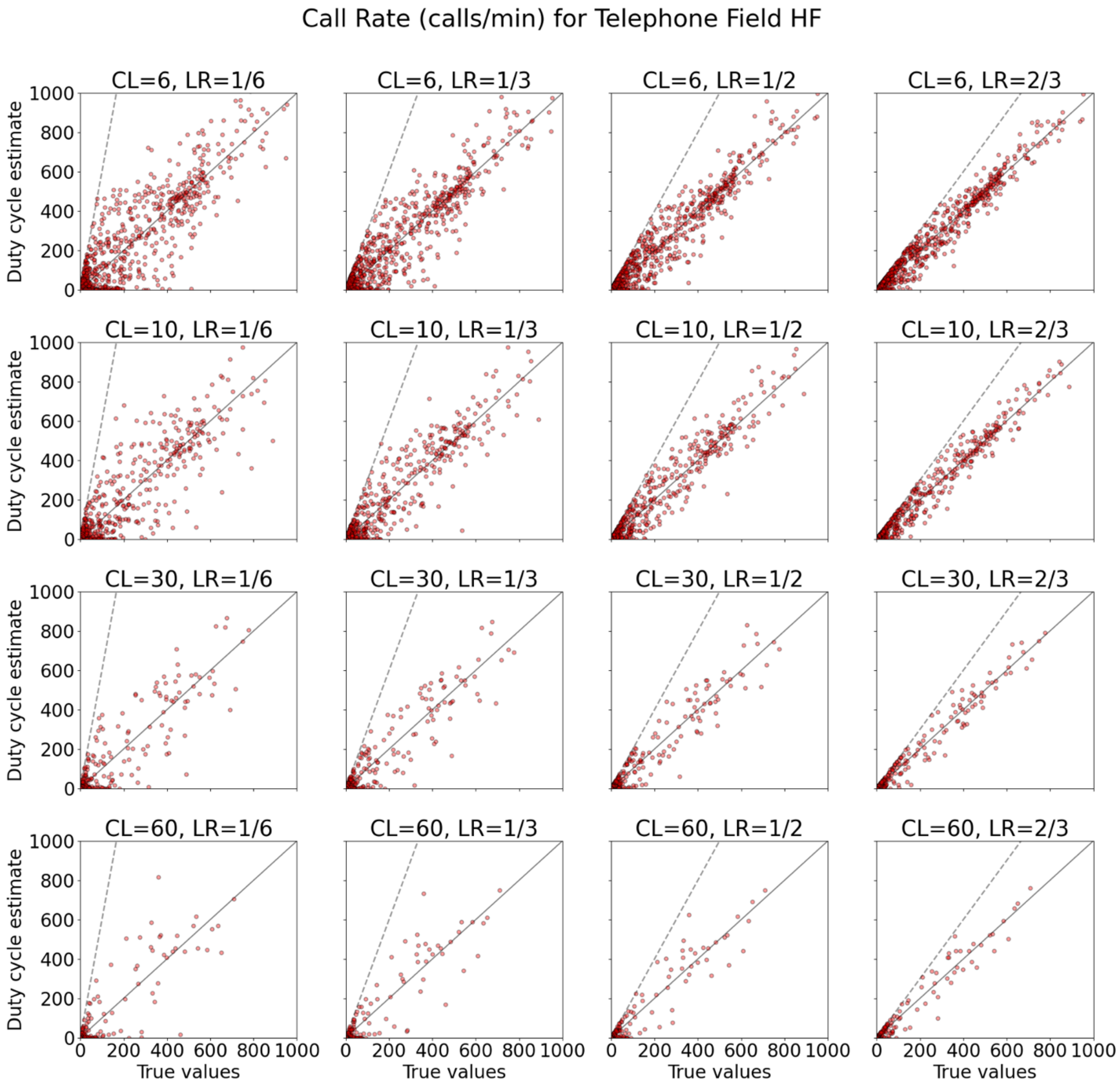


FIG. S9. Scatter plots comparing CR duty-cycle estimates with true CR values across varying duty-cycle schemes from the overall investigation. All other plot details are identical to those in Fig. S6, except all panels here were generated using duty-cycle samples of Telephone Field HF calls during August-September, 2022. **CL**: cycle length (mins); **LR**: listening ratio.

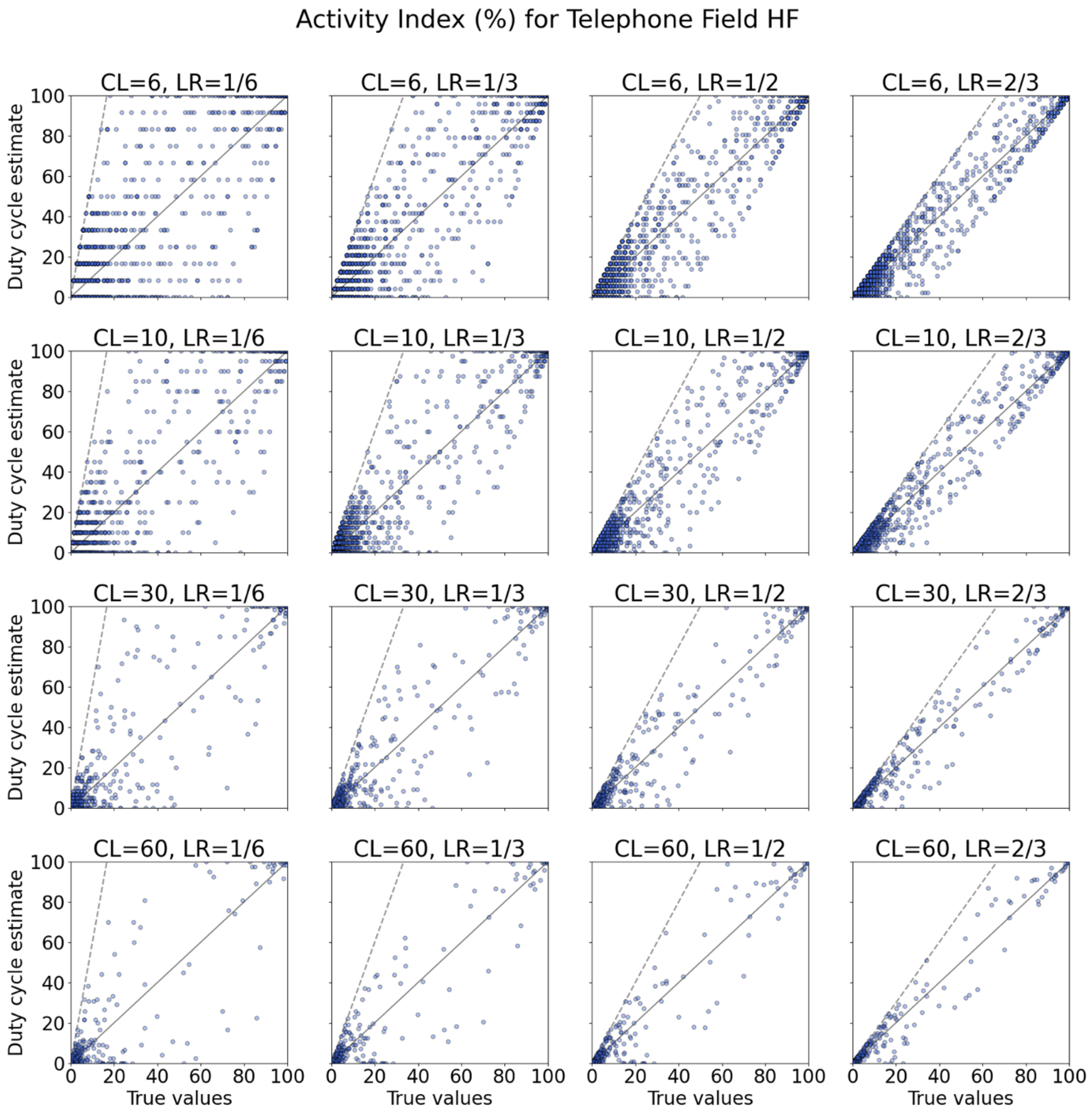


FIG. S10. Scatter plots comparing AI duty-cycle estimates with true AI values across varying duty-cycle schemes from the overall investigation. All other plot details are identical to those in Fig. S9. CL: cycle length (mins); LR: listening ratio.

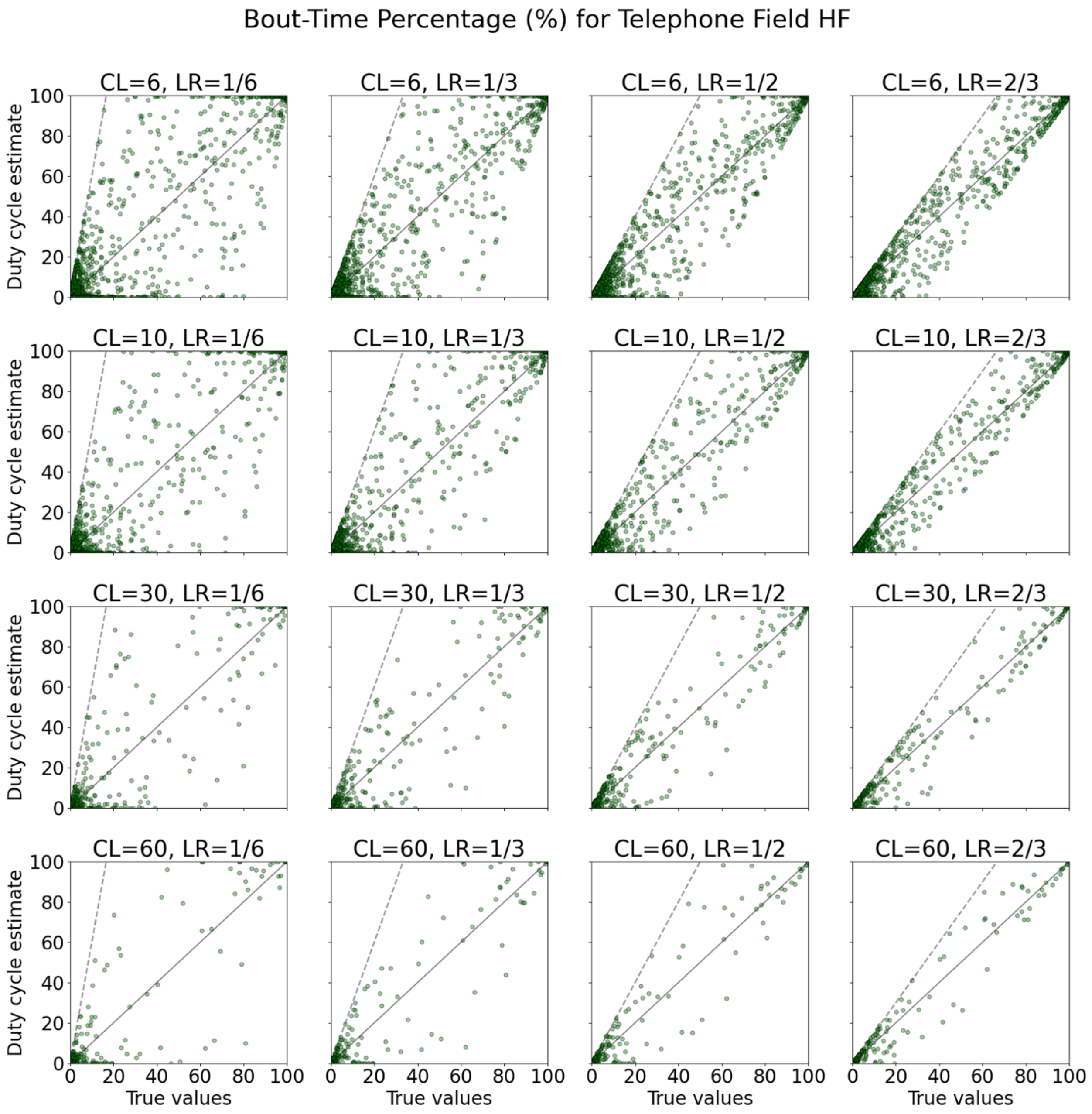


FIG. S11. Scatter plots comparing BTP duty-cycle estimates with true BTP values across varying duty-cycle schemes from the overall investigation. All other plot details are identical to those in Fig. S9. **CL**: cycle length (mins); **LR**: listening ratio.

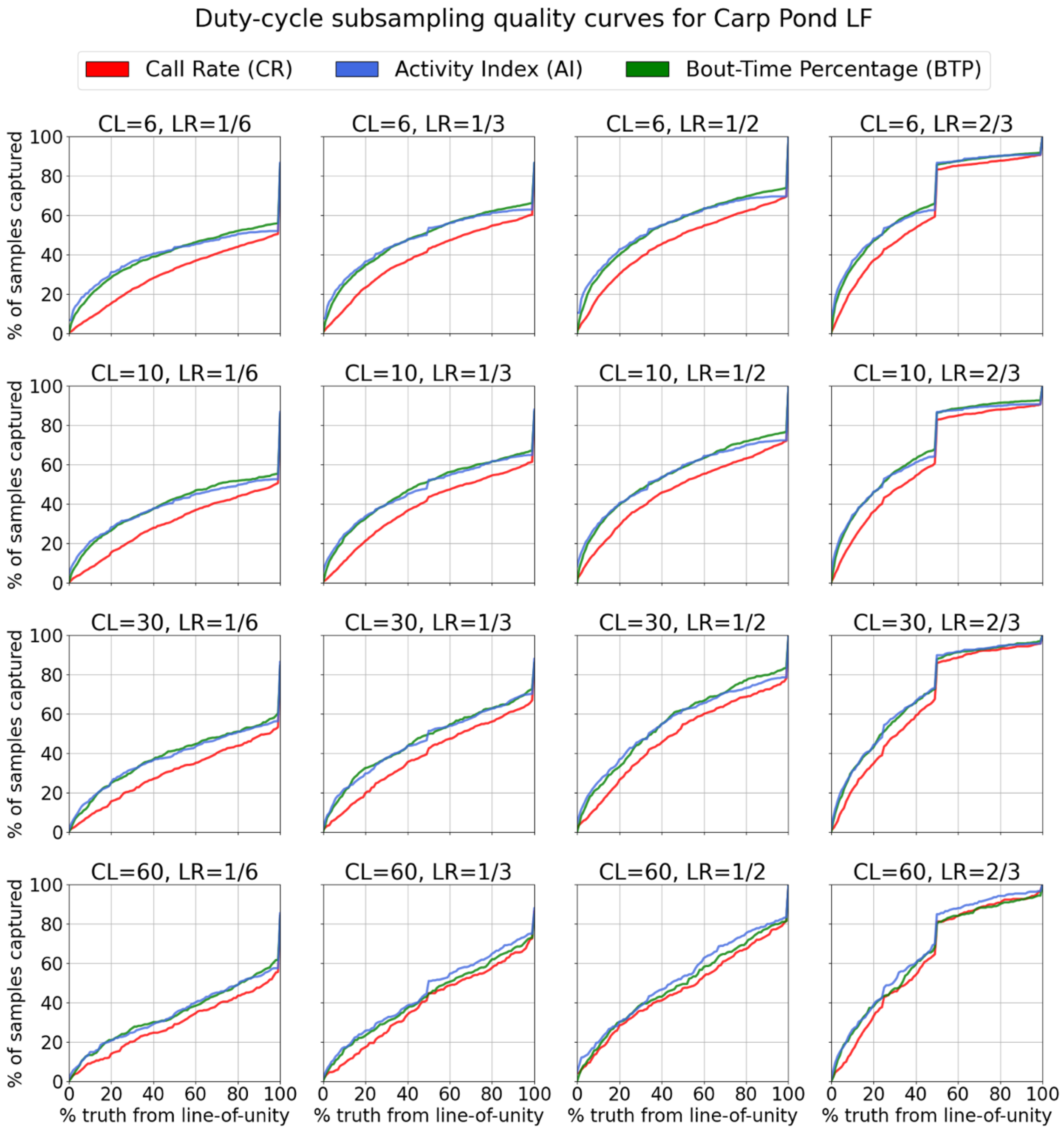


FIG. S12. Plots showing the percentage of cycles meeting the error criterion for varying error bounds across varying duty-cycle schemes from the overall investigation. Each panel in this figure represents the quality of cycle sample estimates under a specific duty-cycle subsampling scheme, denoted by its cycle length (CL; mins) and listening ratio (LR) in the title of the panel. The panel titled "CL: 10, LR: 1/2" is shown as a part of Fig. 4B. Panels with CL of 30min and LRs of ⅙ and ½ are compared in Fig. 6A. Panels with LR of ½ and CLs of 10 min and 60 min are compared in Fig. 6B. Curves that have a faster rise imply a better performance of duty-cycle sample estimation for the specific scheme. All panels were generated using duty-cycle samples of Carp Pond LF calls during July-August, 2022.

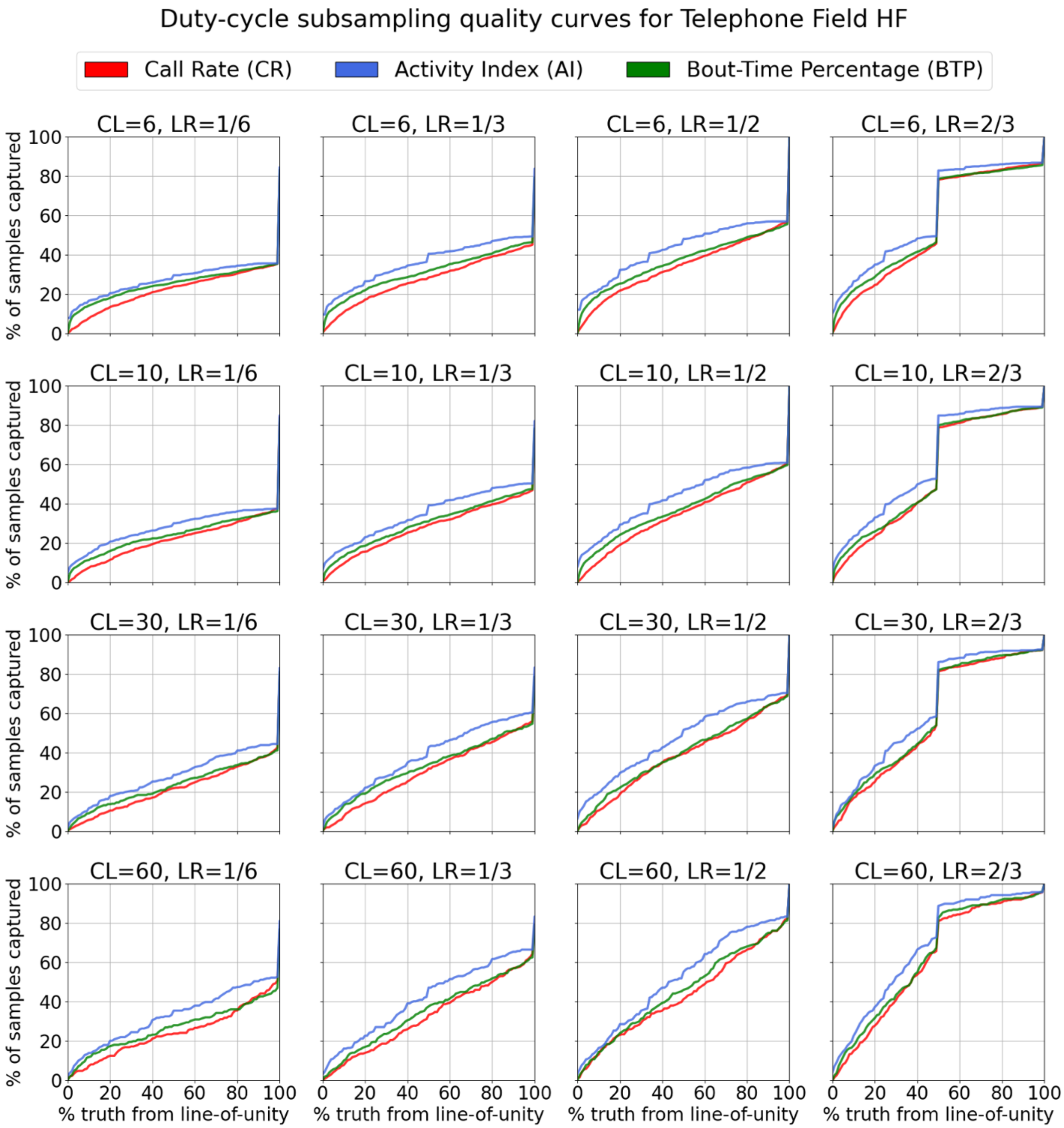


FIG. S13. All plot details are identical to those in Fig. S12, except all panels here were generated using duty-cycle samples of Telephone Field HF calls during August-September, 2022. CL: cycle length (mins); LR: listening ratio.

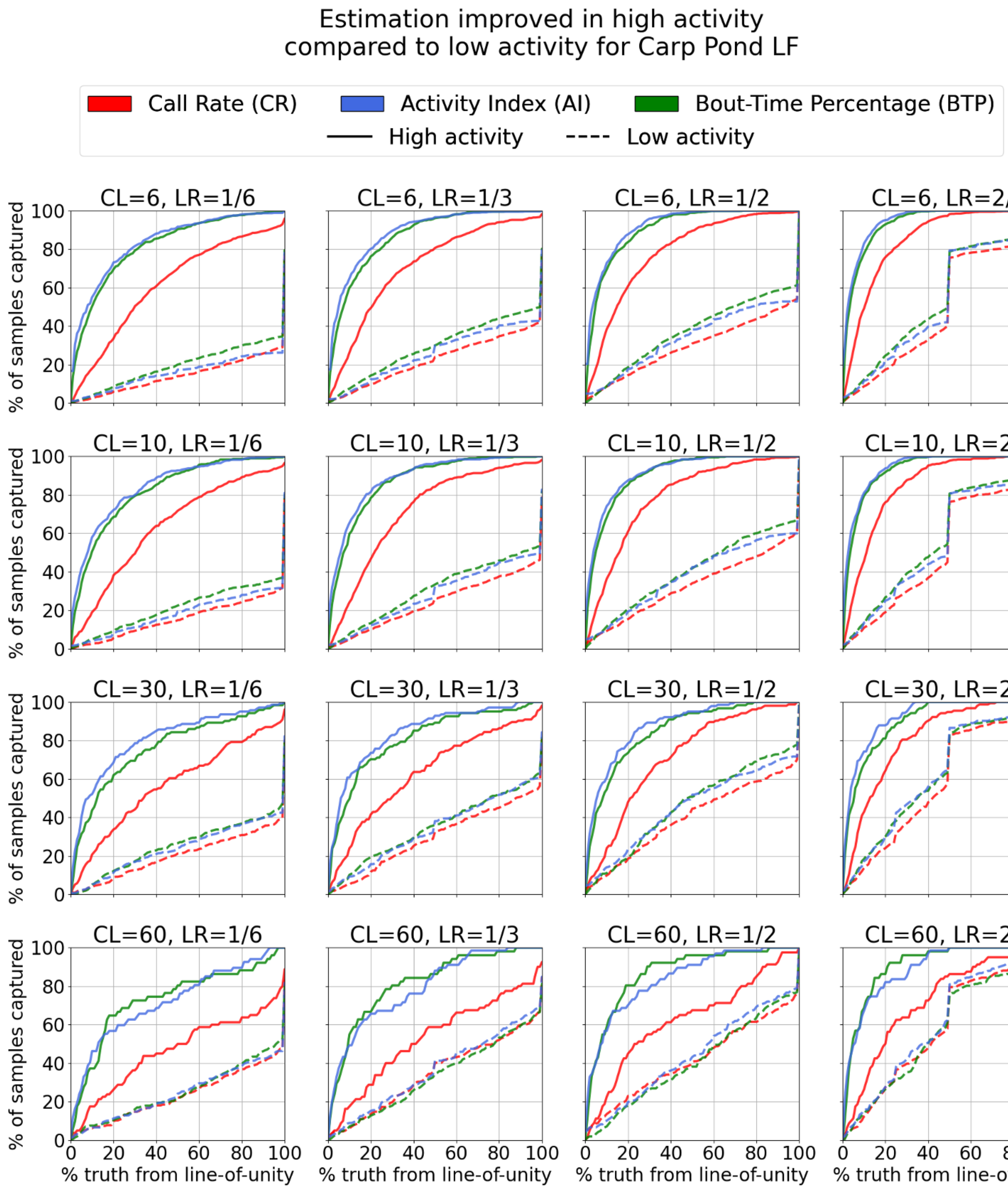


FIG. S14. Plots comparing the influence of duty-cycle subsampling on high and low activity as seen in Fig. 6C. Each panel in this figure represents the comparison of duty-cycle estimation on periods of high and low activity under a specific duty-cycle subsampling scheme, denoted by its cycle length (CL; mins) and listening ratio (LR) in the title of the panel. All sample estimates generated from high activity periods perform better than sample estimates from low activity periods for the same duty-cycle scheme. The panel titled "CL: 10, LR: 1/2" is shown in Fig. 6C. All panels were generated using duty-cycle samples of Carp Pond LF calls during July-August, 2022.

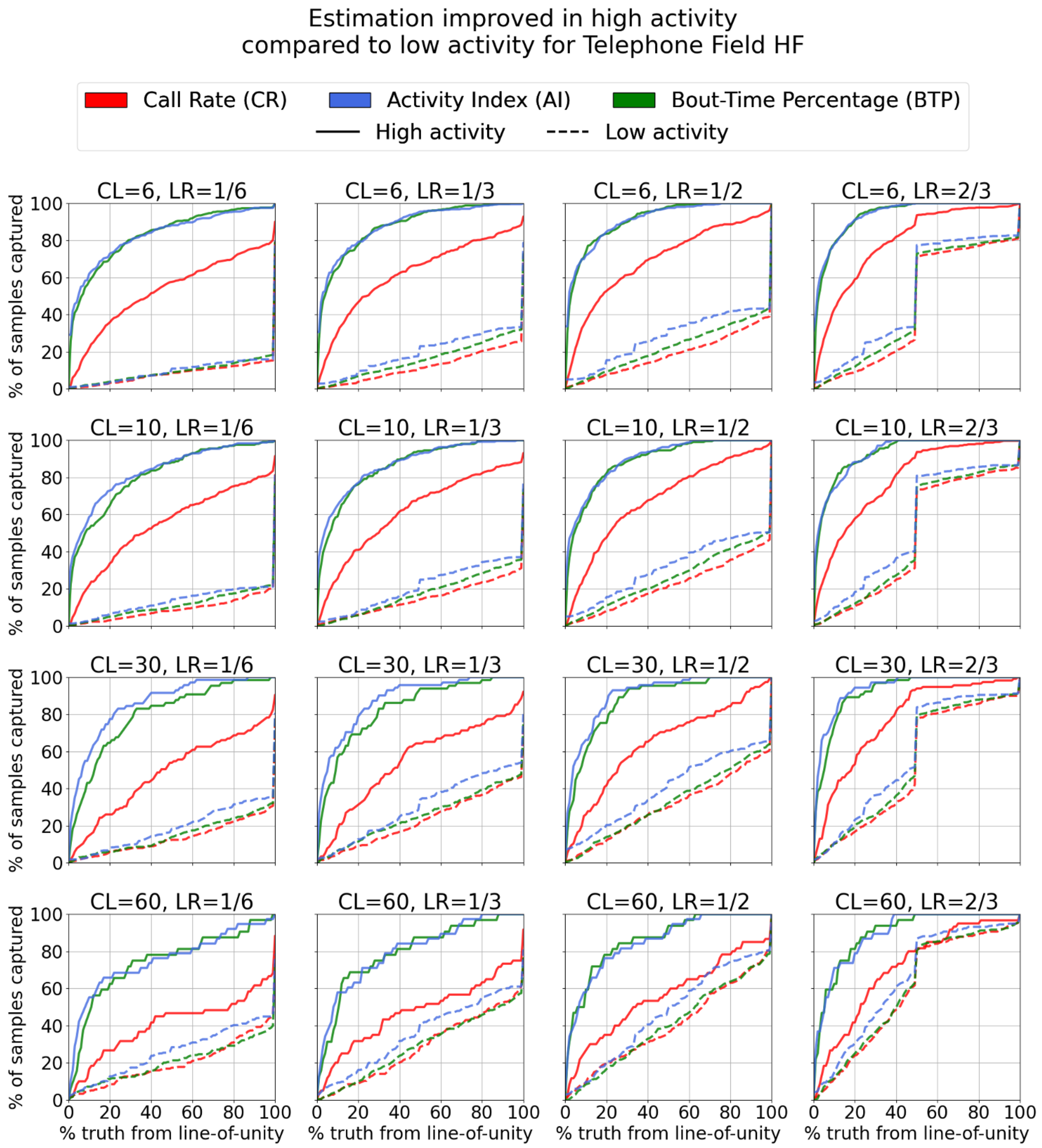


FIG. S15. All plot details are identical to those in Fig. S14, except all panels here were generated using duty-cycle samples of Telephone Field HF calls during August-September, 2022. CL: cycle length (mins); LR: listening ratio.

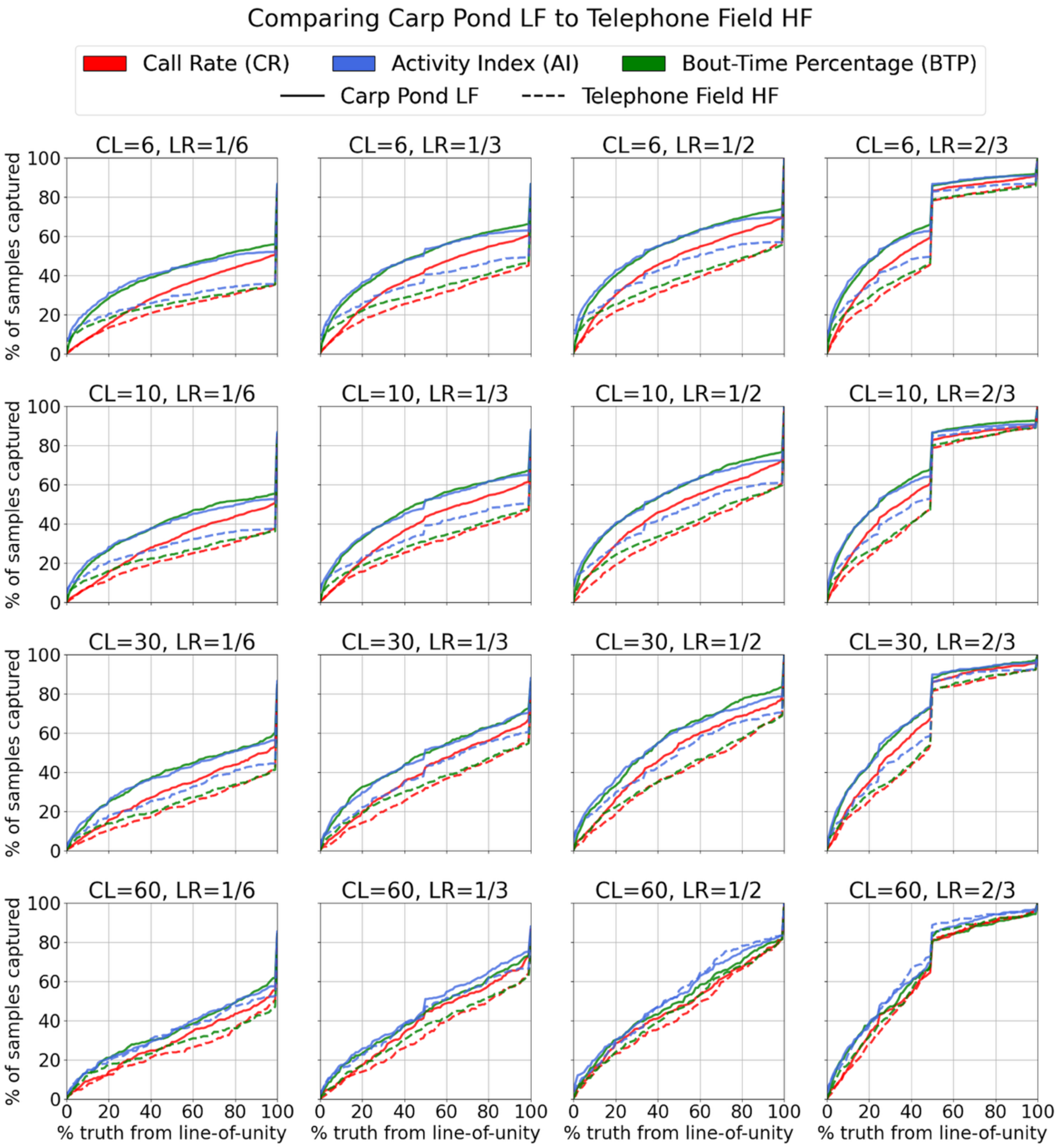


FIG. S16. Plots comparing the influence of duty-cycle subsampling on LF calls and HF calls as seen in Fig. 6D. Each panel in this figure represents the comparison of duty-cycle estimation on periods of LF and HF activity under a specific duty-cycle subsampling scheme, denoted by its cycle length (CL; mins) and listening ratio (LR) in the title of the panel. All sample estimates generated from LF activity perform better than sample estimates from HF activity for the same duty-cycle scheme. The panel titled "CL: 10, LR: 1/2" is shown in Fig. 6D. All panels were generated using duty-cycle samples of Carp Pond LF calls during July-August, 2022 and Telephone Field HF calls during August-September, 2022.
